# The ZmHEAT1/ZmKAKU42–ZmNCH1 module regulates heat stress tolerance by modulating nuclear envelope structure in maize

**DOI:** 10.64898/2026.01.23.701219

**Authors:** Yulong Ji, Shufang Wang, Xiaoliang Wang, Han Liu, Ze Li, Hongmeng Song, Yingjia Han, Wei Chen, Penghao Wu, Xiaohong Yang, Mei Zhang

## Abstract

Heat stress threatens maize production. Through a genome-wide association study using pollen viability decline rate as a phenotype, we identified *ZmHEAT1*, encoding the nuclear envelope protein ZmKAKU41, as a key regulator of thermotolerance. ZmHEAT1 positively regulates heat tolerance at seedling and reproductive stages, with its overexpression improving thermotolerance and its knockout reducing pollen viability and yield under high temperatures. ZmHEAT1 interacts with its homolog ZmKAKU42 and the nucleoskeleton protein ZmNCH1 to form a complex at the nuclear envelope. Mutants lacking ZmHEAT1 or ZmKAKU42 exhibit increased nuclear envelope damage under heat stress, along with repressed expression of key heat shock protein genes (e.g., *HSP90-2*, *HSP17.4*). This work establishes the ZmHEAT1/ZmKAKU42–ZmNCH1 complex as a critical guardian of nuclear integrity and heat response, providing a valuable genetic resource for breeding heat-tolerant maize varieties.

**One-sentence summary:** Modulation of the nuclear envelope structure in maize is involved in heat stress tolerance.

## Introduction

Climate change has led to an increased frequency of extreme weather events, with elevated temperatures causing significant losses in crop production^1^. Maize (*Zea mays*) is particularly sensitive to heat stress during both the seedling and reproductive stages^2^. During the seedling stage, heat stress can lead to membrane damage and excessive accumulation of reactive oxygen species (ROS), which adversely affect development and diminish plant survival rate^3^. During the reproductive stage, heat stress causes a longer interval between anthesis and silking, lowers pollen viability, and potentially leads to male sterility. These factors ultimately contribute to a drop-in seed setting rate and overall lower yields. A recent study revealed that maize inbred lines with smaller tassels may be more susceptible to high temperatures than other inbred lines, such that only lines reaching at least 700 spikelets per tassel are predicted to maintain a consistently high seed setting rate under elevated temperature conditions^4^.

Plants cope with high temperatures through a series of cellular and metabolic responses to sustain growth and development. In particular, the heat stress response (HSR) and the unfolded protein response (UPR) pathways help plants resist high-temperature stress. During initiation of the UPR in maize, the basic leucine zipper protein ZmbZIP60 activates the expression of *HEAT SHOCK FACTOR TRANSCRIPTION FACTOR 13* (*HSFTF13*, also reported as *HSFA6B*), which in turn upregulates the expression of *HEAT UP-REGULATED GENE 1*, encoding an ER chaperone protein, and heat shock protein (HSP) genes under high temperature^5,6^. Intriguingly, ZmbZIP60 appears to have undergone selection during maize domestication and could be associated with the varying degrees of heat tolerance in tropical and temperate maize^7^. In addition, *ZmHSF20*, *ZmHSF4*, *ZmHSFA2*, *ZmCDPK7*, *ZmHSP101*, and *ZmMPK20* were also significantly associated with the heat tolerance of maize^8–12^.

The nucleoskeleton maintains nuclear integrity and chromatin organization at the inner nuclear surface^13^. In Arabidopsis (*Arabidopsis thaliana*), *KAKU4* (Japanese for nucleus) encodes a nuclear envelope protein that is highly expressed in mature pollen; among other effects, loss of KAKU4 function causes reversal of the migration order of the vegetative nuclei and sperm cells in about half of germinated pollen tubes^14,15^. AtKAKU4 localizes at the inner nuclear membrane and interacts with CROWDED NUCLEI 1 (AtCRWN1) and AtCRWN4 to modulate nuclear shape and size^16–18^. The stress-induced localization of nucleoskeleton proteins may be responsible for driving the rearrangement of chromatin inside nuclei. In heat-stressed plants, AtCRWN1 dissociates from the inner nuclear membrane, which could drive nuclear reorganization of the associated chromatin^19^. In maize, CRWNs are represented by NMCP/CRWN HOMOLOG 1 (ZmNCH1) and ZmNCH2, and the KAKU4 homologs are ZmKAKU41 and ZmKAKU42^20^. ZmNCH1, ZmNCH2, and ZmKAKU41 have characteristic properties of linker of nucleoskeleton and cytoskeleton (LINC)-associated plant nuclear envelope proteins and affect overall nuclear architecture. Mutation of *ZmKAKU41* is associated with multiple phenotypes, including changes in nucleus shape in root hairs, development of the stomatal complex, and lower pollen viability^21^. However, how ZmKAKU41 and ZmKAKU42 affect heat tolerance in maize and the underlying functional mechanism remain unknown.

In this study, we performed a genome-wide association study (GWAS) using a panel of 257 maize inbred lines and identified *ZmHEAT1*, encoding the LINC-associated plant nuclear envelope protein ZmKAKU41, as being strongly associated with heat tolerance. ZmHEAT1 forms a complex by interacting with ZmKAKU42 and ZmNCH1 to maintain the structure and integrity of the nuclear envelope and regulate *HSP* genes, affecting the heat tolerance of maize at the seedling and reproductive stages. Field experiments showed that loss of ZmHEAT1 function leads to lower seed setting rate and yield under heat stress conditions. Therefore, ZmHEAT1 represents a potential gene resource for genetic improvement of maize heat tolerance in the future.

## Results

### GWAS for pollen viability decline rate at the reproductive stage in maize

To evaluate the heat tolerance of maize plants at the reproductive stage, we measured the pollen viability of plants maintained under normal conditions or after exposure to high temperature (41°C) during the reproductive stage. We collected pollen grains from a maize population consisting of 257 inbred lines^22^ and calculated the pollen viability decline rate (PVDR), defined as ([pollen viability before treatment − pollen viability after treatment]/pollen viability before treatment) × 100% (Supplementary Figure S1A). We observed substantial variation in PVDR in response to heat stress in this panel, with values ranging from 1.9% (minimal decline in pollen viability) to 99.3% (massive pollen death) (Supplementary Figure S1B). The lines from the tropical/subtropical (TST) subpopulation displayed greater heat tolerance than those from the subpopulations comprising temperate lines (non-stiff stalk) and B73-derived lines (stiff stalk) (Supplementary Figure S1C). Using PVDR as a phenotype, we performed a GWAS, which identified five significant single nucleotide polymorphisms (SNPs) (*p* < 1.03 × 10^−5^) strongly associated with heat tolerance (Figure 1A). Among these, the most significant SNP, chr10.S_146120696 (*p* = 1.51 × 10^−6^), was located within the exon of the gene *ZmHEAT1* (*Zm00001d026487*).

**Fig 1.**
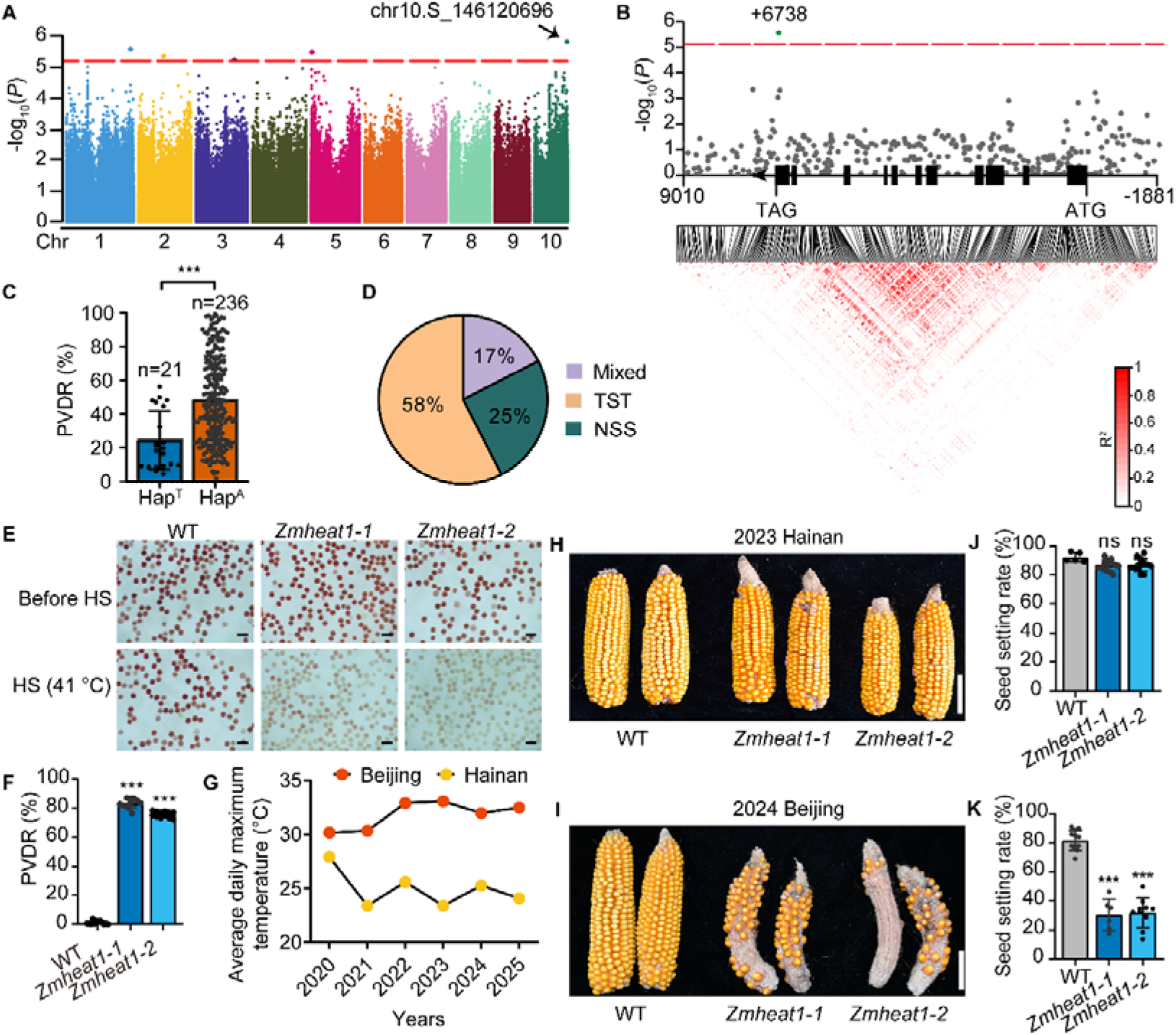
*ZmHEAT1* is associated with maize heat tolerance. A. Manhattan plot of the GWAS results for pollen viability decline rate (PVDR) in the maize diversity panel used in this study. The dashed horizontal line shows the Bonferroni-adjusted significance threshold (*p* = 1.03 × 10^−5^). The leading SNP (chr10.S_146120696) is indicated by an arrow. B. Association analysis of the genetic variation at the *ZmHEAT1* locus and the upstream and downstream 2-kb regions with PVDR in maize and the pattern of pairwise linkage disequilibrium (LD) of polymorphisms. The most significant SNP (+6738) in *ZmHEAT1*. C. PVDR for maize inbred lines sorted by haplotype at the leading SNP. Statistical significance was determined by a two-sided *t*-test (*n* = 21 for haplotype T, *n* = 236 for haplotype A; ***, *p* < 0.001). Values are means ± standard deviation (SD) with all individual data points shown as black dots. D. Distribution of inbred lines with Hap^T^ in different subgroups. TST, tropical/subtropical lines; NSS, temperate non-stiff stalk lines. E. Mature pollen viability of the wild type and gene-edited *Zmheat1* mutants as determined by TTC staining at room temperature. Heat stress (HS) was imposed in an oven at 41□ for 30 min. Scale bars, 100 µm. F. PVDR of the wild type and *Zmheat1* mutants. Statistical significance was determined by one-way ANOVA; ***, *p* < 0.001. Values are means ± SD with all individual data points shown as black dots. G. Average daily maximum temperature profile based on meteorological data for Hainan province and Beijing (2020-2025) at the maize anthesis stage. H, I. Representative photographs of ears from the wild type and the *Zmheat1* mutants grown under normal conditions in 2023 Hainan (H) and heat stress conditions in 2024 Beijing (I). Scale bars, 5 cm. J, K. Seed setting rate of the wild type and *Zmheat1* mutants grown in 2023 Hainan (J) and 2024 Beijing (K). Statistical significance was determined by one-way ANOVA; ns, not significant, ***, *p* < 0.001. Values are means ± SD with all individual data points shown as black dots.

To fully characterize the genetic variation around *ZmHEAT1*, we extracted all SNPs in the *ZmHEAT1* locus and the 2-kb regions upstream and downstream of the locus using high-density genotype data (https://ftp.cngb.org/pub/CNSA/data3/CNP0001565/zeamap/02_Variants/PAN_Zea_Variants/SNP/) for an association analysis of this candidate gene^23^. This revealed the key SNP at position +6738 (A or T; relative to the translational start site, ATG, with the A set as +1) were strongly associated with heat tolerance (Figure 1B). As we did not detect any additional associated SNPs from this analysis, we divided the 257 inbred lines into two haplotypes, Hap^T^ and Hap^A^, based on this SNP. The lines carrying Hap^T^ exhibited a significantly lower PVDR (Figure 1C), suggesting a higher heat tolerance; consistent with this, Hap^T^ lines predominantly belong to the TST subpopulation (Figure 1D). These two haplotypes are related to the substitution of a lysine residue (for the A allele) to a methionine (for the T allele) (Supplementary Figure S2). As *ZmHEAT1* showed constitutive expression (Supplementary Figure S3, A and B), and upregulated expression in different tissues after heat treatment (Supplementary Figure S3, C and D), we analyzed the expression levels of *ZmHEAT1* in the seedlings among 203 lines, and no obvious difference was observed between 20 Hap^T^ and 183 Hap^A^ inbred lines (Supplementary Figure S4). Then, we sought to confirm the difference in heat resilience between Hap^T^ and Hap^A^ haplotypes. We employed a F_2_ population derived from the crosses of maize inbred lines ZZ03 (Hap^T^) and Chang 7-2 (Hap^A^) to conduct phenotypic verification. As a result, the F_2_-*ZmHEAT1*^C72^ displayed a heat-sensitive phenotype after exposure to 45°C for 5 days followed by a 3-day recovery, compared with the F_2_-*ZmHEAT1*^ZZ03^ (Supplementary Figure S5A), with a significantly lower survival rate (Supplementary Figure S5B). The F_2_-*ZmHEAT1*^C72^ also accumulated more ROS than the F_2_-*ZmHEAT1*^ZZ03^ under heat stress but not under normal growth conditions, as determined by DAB staining (Supplementary Figure S5C). Therefore, we considered *ZmHEAT1* the likely candidate gene associated with variation in PVDR, with Hap^T^ being the favorable haplotype resulting in minimal decline of the PVDR following heat stress.

### *ZmHEAT1* positively regulates heat tolerance in maize

To dissect the function of *ZmHEAT1*, we generated *Zmheat1* mutants in the maize inbred line B73-329 through clustered regularly interspaced short palindromic repeats (CRISPR)/CRISPR-associated nuclease 9 (Cas9)-mediated genome editing. We obtained two mutant lines: the *Zmheat1*-*1* mutant contains a 167-bp deletion, and the *Zmheat1-2* mutant contains a 213-bp deletion in the *ZmHEAT1* coding region; both are predicted to lead to early translation termination (Supplementary Figure S2). We assessed the viability of mature pollen from the wild type and the *Zmheat1* knockout lines using 2,3,5-triphenyl tetrazolium chloride (TTC) staining. The pollen viability of *Zmheat1-1* and *Zmheat1-2* plants was lower than that of the wild type specifically under heat stress (Figure 1, E and F).

The seed setting rate of *Zmheat1* mutants was significantly lower than that of wild-type plants when grown in Beijing, China, where high temperatures are frequent at the anthesis stage (Figure 1G). Indeed, the *Zmheat1* mutants produced only a few seeds on each ear, especially the *Zmheat1-2* line (Figure 1, I and K), together with shorter ears (Supplementary Figure S6E), thinner ears (Supplementary Figure S6F), fewer kernels per row (Supplementary Figure S6H), and a lower seed setting rate (Figure 1K), while the number of spike rows was not different from that of the wild type (Supplementary Figure S6G). We observed no significant differences in seed setting rate or yield among genotypes in Hainan, where the climatic conditions are mild (Figure 1, G, H, and J) (Supplementary Figure S6A–D).

At the seedling stage, *ZmHEAT1* expression is induced by heat stress (Supplementary Figure S3, C and D). We thus assessed the function of *ZmHEAT1* during heat stress at the seedling stage. Compared with wild-type seedlings as a control, the two *Zmheat1* mutants displayed a heat-sensitive phenotype after exposure to 45°C for 3 days followed by a 3-day recovery period (Supplementary Figure S7A), with a survival rate of only 10% for the mutants and about 80% for the wild type (Supplementary Figure S7B). The *Zmheat1* mutants accumulated more ROS than the wild type specifically under heat stress conditions, as determined by 3,3′-diaminobenzidine (DAB) staining (Supplementary Figure S7C). Collectively, these data suggest that *ZmHEAT1* regulates heat stress tolerance and yield in maize at the seedling and reproductive stages.

### Overexpression of *ZmHEAT1* improves maize heat tolerance at the seedling and reproductive stages

To validate the role of *ZmHEAT1* in the response to heat stress, we generated an overexpression line for *ZmHEAT1* by driving the *ZmHEAT1* coding sequence from the maize *Ubiquitin* (*Ubi*) promoter in the B73-329 background (*ZmHEAT1*-OE). We chose two OE lines with substantially elevated *ZmHEAT1* transcript levels (Figure 2A), named *ZmHEAT1*-OE #1 and *ZmHEAT1*-OE #2, for analysis. The pollen viability of *ZmHEAT1*-OE #1 and *ZmHEAT1*-OE #2 plants was higher than that of the wild type specifically under heat stress for the plants planted in the greenhouse (Figure 2, B and C). Further, we analyzed the seed setting rates and ear traits of wild-type and *ZmHEAT1*-OE plants planted in Hainan and Beijing field, separately. As a result, the seed setting rate of wild-type was significantly lower than that of *ZmHEAT1*-OE plants when grown in Beijing (Figure 2, F and G), while no obvious differences were observed when grown in Hainan (Figure 2, D and E; Supplementary Figure S8A–D) and no obvious differences in any ear traits among genotypes in Beijing and Hainan (Supplementary Figure S8E–H).

**Fig 2.**
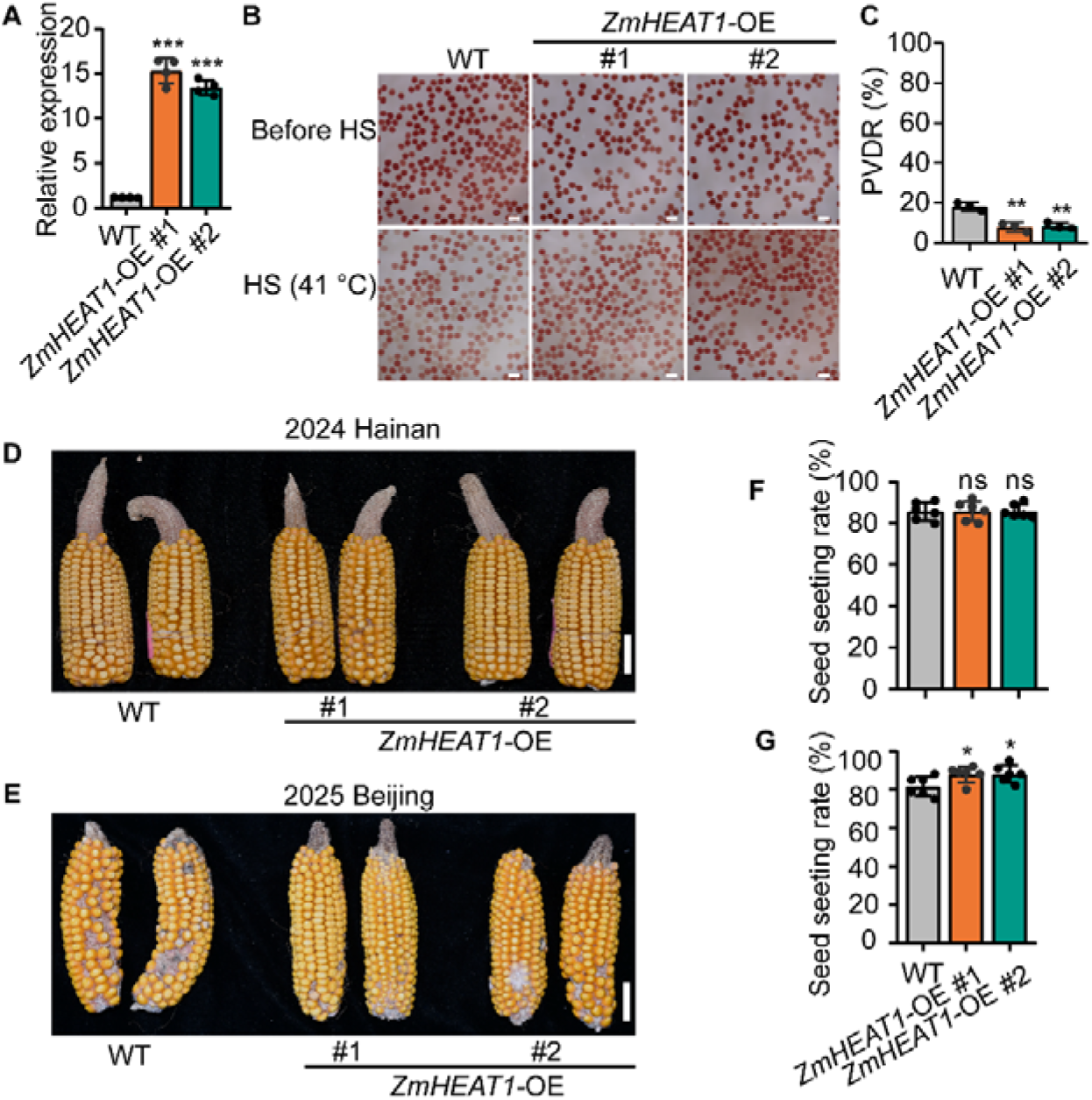
*ZmHEAT1* improves the heat tolerance of maize. A. Relative *ZmHEAT1* expression levels in the wild type and *ZmHEAT1-*OE #1 and *ZmHEAT1*-OE #2. Statistical significance was determined by one-way ANOVA; ***, *p* < 0.001. Values are means ± SD with all individual data points shown as black dots. B. Analysis of mature pollen viability for wild-type and *ZmHEAT1-*OE plants as determined by TTC staining at room temperature. Heat stress (HS) was imposed in an oven at 41°C for 30 min. Scale bars, 100 µm. C. Pollen viability decline rate (PVDR) of wild-type and *ZmHEAT1-*OE plants. Statistical significance was determined by one-way ANOVA; **, *p* < 0.01. Values are means ± SD with all individual data points shown as black dots. D, E. Representative photographs of ears from the wild type and the *ZmHEAT1*-OE mutants grown under normal conditions in 2024 Hainan (D) and heat stress conditions in 2025 Beijing (E). Scale bars, 5 cm. F, G. Seed setting rate of the wild type and *ZmHEAT1*-OE mutants grown in 2024 Hainan (F) and in 2025 Beijing (G). Statistical significance was determined by one-way ANOVA; ns, not significant, *, *p* < 0.05. Values are means ± SD with all individual data points shown as black dots.

At the seedling stage, both *ZmHEAT1*-OE #1 and *ZmHEAT1*-OE #2 lines showed a heat-tolerant phenotype compared with the wild type, as evidenced by minimal leaf wilting and death (Supplementary Figure S9A), together with greater survival rates (Supplementary Figure S9B) and more ROS accumulation (Supplementary Figure S9C). Collectively, these data suggest that overexpression of *ZmHEAT1* improves heat tolerance in maize at seedling and reproductive stages.

### ZmHEAT1 and ZmKAKU42 localize at the nuclear envelope and interact with each other

Phylogenetic analysis indicates that *ZmHEAT1* is an ortholog of Arabidopsis *KAKU4* (Supplementary Figure S10A), a well-characterized nuclear envelope protein involved in nuclear morphology and stress response^13–16,18^. In maize, *ZmHEAT1* has a paralog gene, *ZmKAKU42* (Supplementary Figure S10, A and B). We wondered whether the nuclear envelope takes on an alternative organization under heat stress; accordingly, we examined the localization of *Z*mHEAT1 and ZmKAKU42 in plants grown under normal temperature conditions and after heat stress. To this end, we infiltrated constructs encoding fusions between ZmHEAT1 or ZmKAKU42 and green fluorescent protein (GFP) into the leaves of *N. benthamiana* plants. We detected GFP fluorescence signals for ZmHEAT1-GFP and ZmKAKU42-GFP at the nuclear envelope regardless of exposure to heat stress (Supplementary Figures S11 and S12). To investigate the relationship between ZmHEAT1 and ZmKAKU42, we conducted a yeast two-hybrid assay, which revealed that ZmHEAT1 interacts with ZmKAKU42 (Figure 3A). We also performed a glutathione S-transferase (GST) in vitro pull-down assay by mixing recombinant purified ZmKAKU42-maltose-binding protein (MBP) with ZmHEAT1-GST or GST alone. SDS-PAGE and immunoblot analysis of the proteins bound to glutathione beads detected ZmKAKU42-MBP among the proteins pulled down by ZmHEAT1-GST, but not by GST (Figure 3B), suggesting that ZmHEAT1 interacts with ZmKAKU42 in vitro.

**Fig 3.**
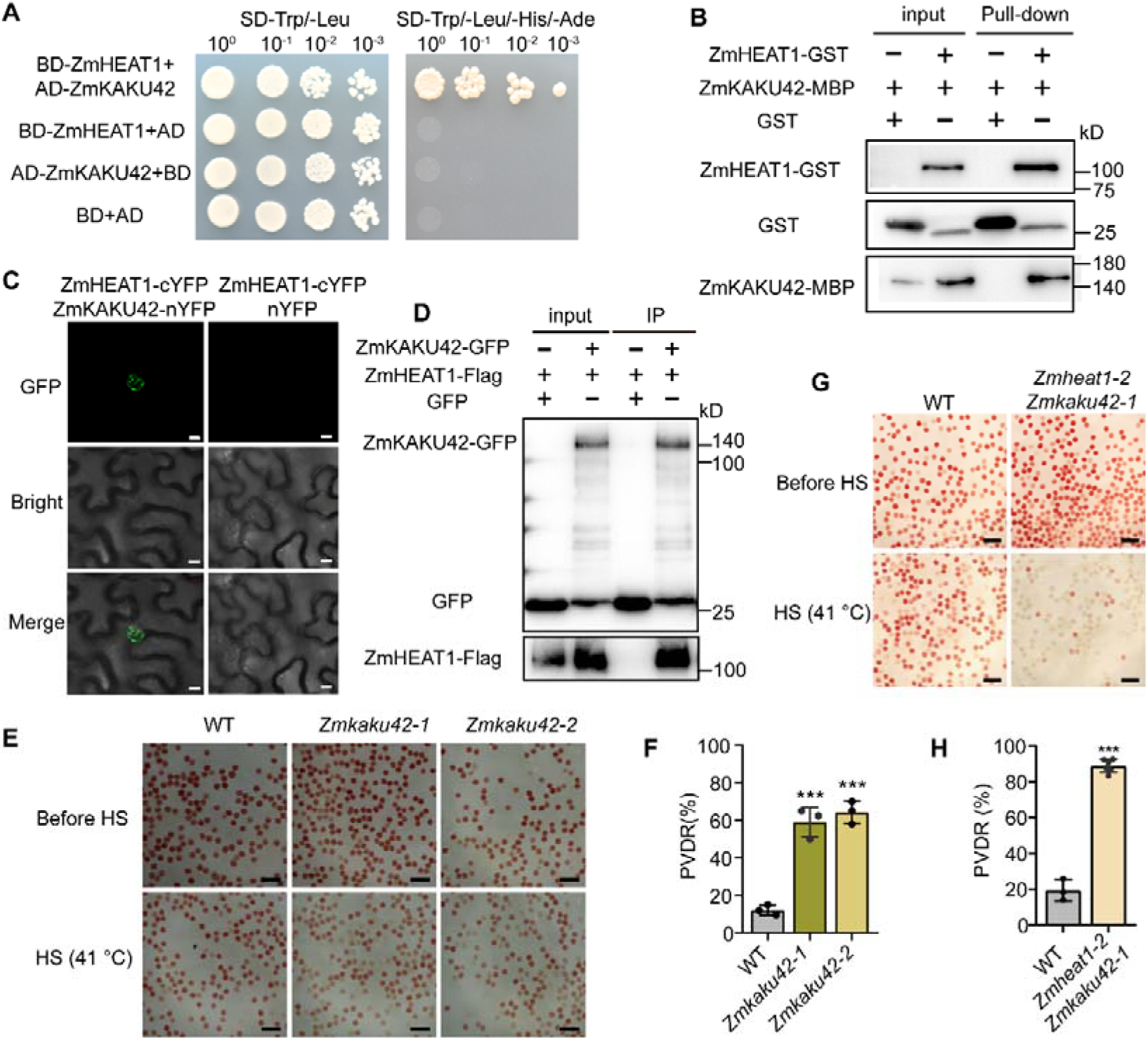
ZmHEAT1 interacts with ZmKAKU42, and ZmKAKU42 improves the heat tolerance of maize. A. Yeast two-hybrid (Y2H) assay showing that ZmHEAT1 interacts with ZmKAKU42. Yeast cells were grown on synthetic defined (SD) medium lacking Trp and Leu (−Trp−Leu) or Trp, Leu, His, and Ade (−Trp −Leu −His −Ade). B. In vitro glutathione S-transferase (GST) pull-down assay showing the interaction between recombinant purified ZmHEAT1-GST and ZmKAKU42-MBP. Recombinant purified GST or ZmHEAT1-GST was immobilized on glutathione agarose beads and incubated with ZmKAKU42-MBP. ZmKAKU42 was detected with an anti-MBP antibody. GST was used as a negative control. C. Bimolecular fluorescence complementation (BiFC) assay showing the interaction of ZmHEAT1 with ZmKAKU42 in *N. benthamiana* leaf cells. The YFP signal was visualized by confocal microscopy. Scale bars, 10 µm. D. Co-immunoprecipitation (Co-IP) assay showing the interaction between ZmHEAT1 and ZmKAKU42 in protein extracts from *N. benthamiana* leaves expressing the indicated constructs. Input and co-immunoprecipitated proteins were analyzed using anti-GFP and anti-Flag antibodies. GFP was used as a negative control. E. Analysis of mature pollen viability of the wild type and *Zmkaku42* mutants as determined by TTC staining at room temperature. Heat stress (HS) was imposed in an oven at 41□ for 30 min. Scale bars, 100 µm. F. PVDR of the wild type and *Zmkaku42* mutants. Statistical significance was determined by one-way ANOVA; ***, *p* < 0.001. Values are means ± SD with all individual data points shown as black dots. G. Analysis of mature pollen viability of the wild type and *Zmheat1 Zmkaku42* mutants as determined by TTC staining at room temperature. HS was imposed in an oven at 41□ for 30 min. Scale bars, 100 µm. H. PVDR of the wild type and *Zmheat1 Zmkaku42* mutants. Statistical significance was determined by one-way ANOVA; ***, *p* < 0.001. Values are means ± SD with all individual data points shown as black dots.

To confirm the interaction between ZmHEAT1 and ZmKAKU42 in planta, we performed a bimolecular fluorescence complementation (BiFC) assay in *N. benthamiana*. We detected yellow fluorescent protein (YFP) in the nuclei of epidermal cells following co-infiltration with *ZmHEAT1-cYFP* (encoding a fusion of ZmHEAT1 with the C-terminal half of YFP) and *ZmKAKU42-nYFP* (encoding a fusion of ZmKAKU42 with the N-terminal half of YFP) constructs, but not when *ZmHEAT1-cYFP* was co-infiltrated with *nYFP* (Figure 3C). We also performed a coimmunoprecipitation (Co-IP) assay with an anti-GFP antibody using total protein extracts from *N. benthamiana* leaves co-infiltrated with ZmHEAT1-Flag and ZmKAKU42-GFP constructs. We detected ZmHEAT1-Flag among the proteins co-precipitated with ZmKAKU42-GFP, but not those with GFP alone (Figure 3D). Therefore, ZmHEAT1 interacts with ZmKAKU42 in vitro and in vivo.

To determine the function of ZmKAKU42, we generated *Zmkaku42* mutants in the B73-329 inbred line through CRISPR/Cas9-mediated genome editing. We obtained two mutant lines: the *Zmkaku42*-1 mutant contains a 113-bp deletion (premature termination of protein coding), and the *Zmkaku42-2* mutant contains a 7-bp deletion and a 1-bp insertion in the *ZmKAKU42* coding region (predicted to produce a truncated form of ZmKAKU42) (Supplementary Figure S13 and S14). When we assessed the viability of mature pollen from the wild type and the two *Zmkaku42* knockout lines using TTC staining, we discovered that pollen viability is much lower in the *Zmkaku42-1* and *Zmkaku42-2* mutants compared with the wild type under heat stress (Figure 3, E and F). At the seedling stage, the two *Zmkaku42* mutants displayed a heat-sensitive phenotype after exposure to 45°C for 3 days followed by a 3-day recovery, compared with the wild type (Supplementary Figure S15A), with a significantly lower survival rate (Supplementary Figure S15B). The two *Zmkaku42* mutants also accumulated more ROS than the wild type under heat stress but not under normal growth conditions, as determined by DAB staining (Supplementary Figure S15C). To further decipher the relationship between *ZmHEAT1* and *ZmKAKU42*, we generated the *Zmheat1-2 Zmkaku42-1* double mutant by crossing *Zmheat1-2* to *Zmkaku42-1*. When we assessed the viability of mature pollen from the wild type and the *Zmheat1-2 Zmkaku42-1* double mutant using TTC staining, we discovered that pollen viability is much lower in the *Zmheat1-2 Zmkaku42-1* mutants compared with the wild type under heat stress (Figure 3, G and H). Meanwhile, the *Zmheat1-2 Zmkaku42-1* mutants displayed a heat-sensitive phenotype after exposure to 45°C for 3 days followed by a 3-day recovery, compared with the wild type (Supplementary Figure S16A), with a significantly lower survival rate (Supplementary Figure S16B). The *Zmheat1-2 Zmkaku42-1* mutants also accumulated more ROS than the wild type under heat stress but not under normal growth conditions, as determined by DAB staining (Supplementary Figure S16C). Together, these data suggest that ZmHEAT1 and ZmKAKU42 collectively contribute to heat stress tolerance in maize at the seedling and reproductive stages.

### ZmHEAT1 and ZmKAKU42 interact with ZmNCH1 at the nuclear envelope

In Arabidopsis, KAKU4 physically interacts with the nuclear matrix proteins CRWN (CRWN1 and CRWN4)^16,18^. In maize, ZmNCH1 and ZmNCH2 as the orthologs of AtCRWNs^20^. Therefore, we wondered whether ZmNCH1 and/or ZmNCH2 interacts with ZmHEAT1 and/or ZmKAKU42 in maize. To investigate the relationship of ZmHEAT1/ZmKAKU42 and ZmNCH1/ZmNCH2, we first investigated the localization of ZmNCH1-GFP and ZmNCH2-GFP in *N. benthamiana*. We detected the two fusion proteins in the nucleus regardless of exposure to heat stress conditions (Supplementary Figures S17 and S18). Then, we conducted a yeast two-hybrid assay, which revealed the interaction of ZmHEAT1 and ZmKAKU42 with ZmNCH1 (Figure 4A), whereas ZmNCH2 failed to interact with ZmHEAT1 or ZmKAKU42 (Supplementary Figure S19, A and B). We performed a GST in vitro pull-down assay by mixing recombinant purified ZmHEAT1-GST, ZmKAKU42-GST, or GST with recombinant purified ZmNCH1-His. Immunoblot analysis of the proteins bound to the glutathione beads showed that ZmNCH1-His is pulled down by ZmHEAT1-GST and ZmKAKU42-GST, but not by GST (Figure 4, B and C), suggesting that ZmHEAT1 and ZmKAKU42 interact with ZmNCH1 in vitro.

**Fig 4.**
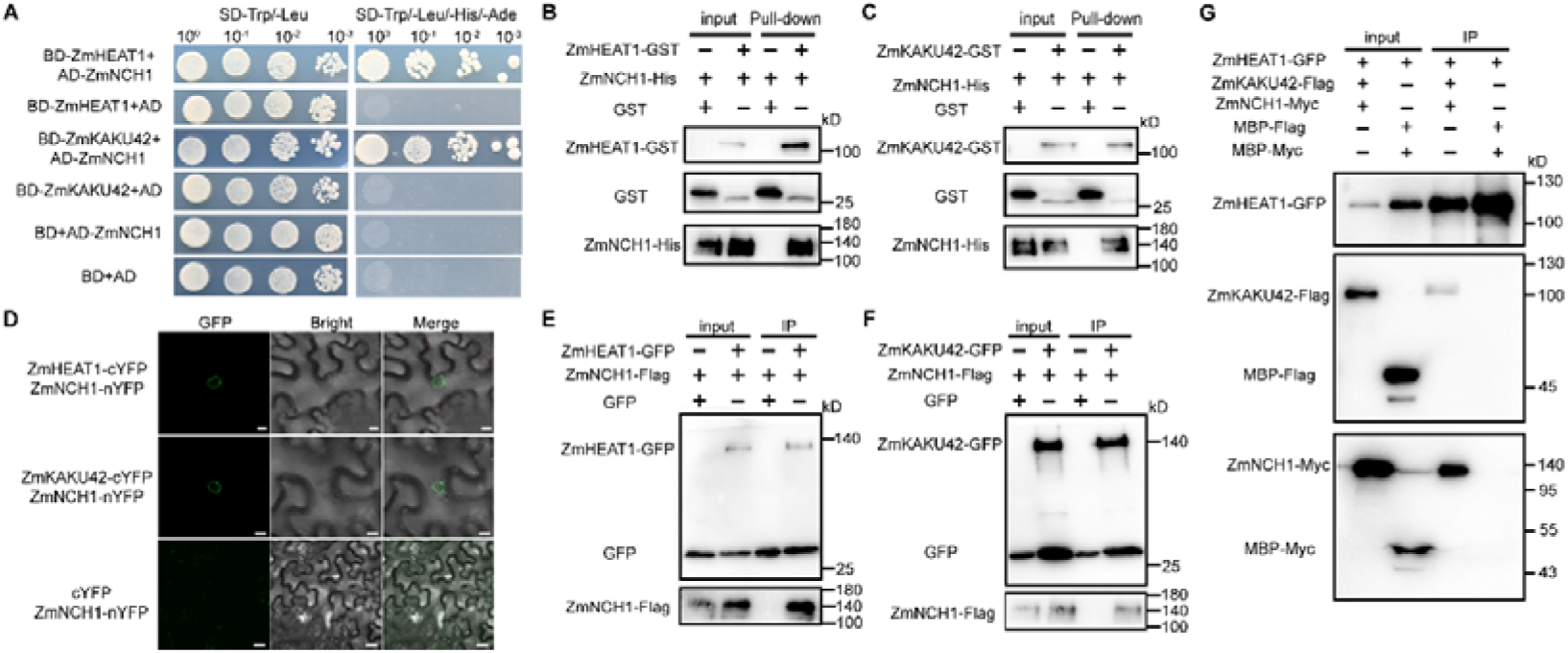
ZmHEAT1 and ZmKAKU42 interact with ZmNCH1. A. Y2H assays showing that ZmHEAT1 and ZmKAKU42 interact with ZmNCH1. Yeast cells were grown on SD medium lacking Trp and Leu (−Trp −Leu) or Trp, Leu, His, and Ade (−Trp −Leu −His −Ade). B, C In vitro GST pull-down assay showing that recombinant purified ZmHEAT1 (B) and ZmKAKU42 (C) interact with ZmNCH1. Recombinant purified GST, ZmHEAT1-GST, or ZmKAKU42-GST was immobilized on glutathione agarose beads and incubated with recombinant purified ZmNCH1-His. ZmNCH1 was detected with an anti-His antibody. GST was used as a negative control. D. BiFC assay showing that ZmHEAT1 and ZmKAKU42 interact with ZmNCH1 in *N. benthamiana* leaf cells. The YFP signal was visualized by confocal microscopy. Scale bars, 10 µm. E, F. Co-IP assay showing the interaction between ZmHEAT1 (E) or ZmKAKU42 (F) and ZmNCH1 in *N. benthamiana* leaves. Input and co-immunoprecipitated proteins were detected using anti-GFP and anti-Flag antibodies. GFP was used as a negative control. G. Co-IP assay showing the interaction between ZmHEAT1, ZmKAKU42 and ZmNCH1 in protein extracts from *N. benthamiana* leaves expressing the indicated constructs. Input and co-immunoprecipitated proteins were analyzed using anti-GFP, anti-Flag and anti-Myc antibodies. MBP-Flag and MBP-Myc was used as a negative control.

To confirm the interaction between ZmHEAT1, ZmKAKU42, and ZmNCH1 in planta, we performed a BiFC assay in *N. benthamiana*. We observed the reconstitution of YFP in the nuclei of epidermal cells when co-infiltrated with *ZmHEAT1-cYFP* or *ZmKAKU42-cYFP* and *ZmNCH1-nYFP* constructs (Figure 4D), but not when *ZmNCH1-nYFP* was co-infiltrated with *cYFP*. We also performed a Co-IP assay with an anti-GFP antibody using total protein extracts from *N. benthamiana* leaves co-infiltrated with ZmHEAT1-GFP or ZmKAKU42-GFP and ZmNCH1-Flag constructs. We detected ZmNCH1-Flag among the proteins immunoprecipitated with ZmHEAT1-GFP or ZmKAKU42-GFP, but not with GFP alone (Figure 4, E and F). To determine the function of *ZmNCH1*, we also generated *ZmNCH1*-OE lines in Arabidopsis. We treated 7 days Arabidopsis seedlings at 43°C for 1.5 hours, and after 7 days of recovery under normal temperature conditions, the survival rate of *ZmNCH1*-OE plants was significantly higher than WT plants (Supplementary Figure S20, A and B).

To further verify whether ZmHEAT1/ZmKAKU42 and ZmNCH1 form a complex, we performed a Co-IP assay using an anti-GFP antibody with total protein extracts from *N. benthamiana* leaves co-infiltrated with ZmHEAT1-GFP, ZmKAKU42-Flag, and ZmNCH1-Myc constructs. As a negative control, leaves co-expressing ZmHEAT1-GFP together with MBP-Flag and MBP-Myc fusion proteins (MBP tag added to facilitate molecular□weight discrimination) were included. Both ZmKAKU42-Flag and ZmNCH1-Myc were specifically co□precipitated with ZmHEAT1□GFP, whereas neither MBP-Flag nor MBP-Myc was detected (Figure 4G). Meanwhile, we assessed the effect of heat stress on these protein-protein interactions in vivo using a quantitative bimolecular fluorescence complementation (BiFC) assay. Fluorescence intensity was measured in tobacco cells co-expressing ZmHEAT1 and ZmKAKU42, or ZmHEAT1, ZmKAKU42, and ZmNCH1, under control and heat stress conditions (Supplementary Figure S21A). Statistical analysis showed that the interaction strength in both configurations was significantly reduced upon heat treatment (Supplementary Figure S21, A-D). These data indicate that the formation/stability of the ZmHEAT1□containing complexes is negatively regulated by elevated temperature. Therefore, ZmHEAT1 and ZmKAKU42 with ZmNCH1 form a complex in vitro and in vivo.

### The ZmHEAT1/ZmKAKU42–ZmNCH1 complex regulates nuclear envelope structure and *HSP* expression

To explore the effect of the ZmHEAT1/ZmKAKU42–ZmNCH1 complex on the nuclear envelope, we subjected 14-day-old seedlings to high-temperature treatment and observed nuclei in their leaves under transmission electron microscopy before and after treatment. We did not observe a substantial difference in the nuclear envelope between the wild type and *Zmheat1-2*, *Zmkaku42-1*, *Zmheat1-2 Zmkaku42-1* and *ZmHEAT1-*OE#1 under control conditions (Figure 5A). However, following 48 h of heat stress, the *Zmheat1-2*, *Zmkaku42-1*, and the *Zmheat1-2 Zmkaku42-1* mutants displayed a significant increase in the frequency of nuclear envelope damage (Figure 5, A and B). In contrast, *ZmHEAT1*-OE lines showed a clear reduction in damage incidence, supporting a protective role of *ZmHEAT1* under prolonged stress (Figure 5, A and B).

**Fig 5.**
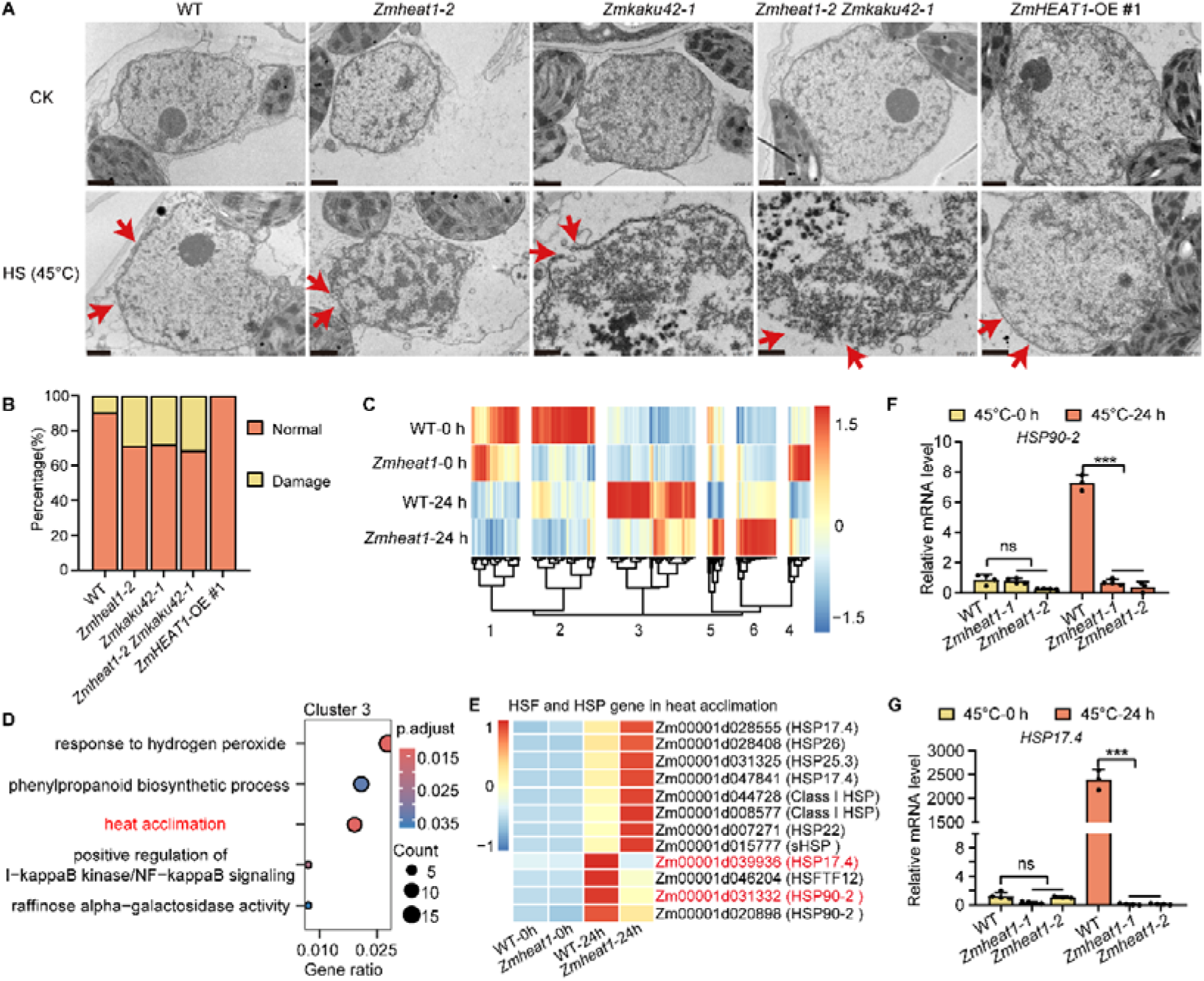
Transcriptome analysis of ZmHEAT1-dependent heat-regulated genes. A. Transmission electron microscopy (TEM) analysis of the structure of the nuclear envelope in the wild type, *Zmheat1-2*, *Zmkaku42-1*, *Zmheat1-2 Zmkaku42-1* and the *ZmHEAT1*-OE #1 mutant under normal (CK) or HS conditions. The leaves were collected from V3-stage seedlings grown at 28°C/22°C (CK) alone or exposed to 45°C for 48 h. Scale bars, 1 µm. The red arrows indicate the nuclear envelope. B. Nuclear envelope damage percentage in WT, *Zmheat1-2*, *Zmkaku42-1*, *Zmheat1-2 Zmkaku42-1* and the *ZmHEAT1*-OE #1 mutants exposed to 45°C for 48 h. C. Heatmap representation of gene expression levels following clustering analysis of ZmHEAT1-dependent heat-regulated genes (HTGs). Heatmap shows log_2_-normalized (TPM) values of 1,812 DEGs. DEGs were clustered with the R package Pheatmap based on their expression patterns. D. Gene Ontology (GO) term enrichment analysis of the DEGs from cluster 3. The size of the circles indicates the number of DEGs for each GO term, while their colors represent the associated adjusted *p*-value. E. Heatmap of gene expression for HSF and HSP related genes in Cluster 3 under heat acclimation. F, G. Relative expression levels for the *HSP* genes *HSP90-2* (*Zm00001d031332*) (E) and *HSP17.4* (*Zm00001d039936*) (F) in wild-type and *Zmheat1* mutant seedlings before and after 45°C treatment, as determined by RT-qPCR. Expression in the wild type before treatment was set to 1. Statistical significance was determined by one-way ANOVA; ns, not significant, ***, *p* < 0.001. Values are means ± SD with all individual data points shown as black dots.

We performed transcriptome deep sequencing (RNA-seq) from 14-day-old seedlings to explore the expression of downstream genes affected by ZmHEAT1. We identified differentially expressed genes (DEGs) based on the criteria of a significant difference (*p* < 0.05) with an absolute log_2_(fold-change) ≥ 1. We identified 9,793 DEGs in wild-type seedlings after exposure to heat (45°C-24 h) compared with wild-type seedlings before heat treatment (45°C-0 h), among which 4,933 were upregulated and 4,860 were downregulated (Supplementary Figure S22A, S23, Figure 5C, Supplementary Data Set S1). To identify ZmHEAT1-regulated DEGs, we defined the DEGs in a comparison between *Zmheat1* seedlings with or without heat treatment (45°C) for 24 h, leading to a set of 3,812 DEGs regulated by ZmHEAT1; we also identified 2,233 DEGs from the comparison of *Zmheat1* versus WT under normal conditions and 2,629 DEGs from the comparison of *Zmheat1* versus WT after 24 h of heat treatment (Supplementary Figure S22, B and C, S23, Supplementary Data Sets S2 and S3). Of the 9,793 DEGs identified in the wild type in response to heat stress treatment and the 3,812 DEGs identified in *Zmheat1* upon exposure to heat stress, 1,812 DEGs were shared, defining genes that are regulated by both heat and ZmHEAT1 (Supplementary Figure S23 and Supplementary Data Set S4).

Clustering analysis grouped the 1,812 ZmHEAT1-dependent heat-regulated genes (HTGs) into six clusters (clusters 1–6) (Figure 5C). Based on the expression patterns of the genes from each of the six clusters, we focused on cluster 3, a class of genes that showed significantly altered expression in the *Zmheat1* mutant compared to the WT in response to heat stress. We performed a Gene Ontology (GO) term enrichment analysis for the genes of cluster 3 to explore their associated biological processes. Cluster 3 was enriched in genes related to biological processes involved in the response to hydrogen peroxide, phenylpropanoid biosynthetic process, and heat acclimation (Figure 5D). Many different *HSP* and *HSF* genes were enriched in these terms, including *HSFTF12*, *HSP90-2*, and *HSP17.4* (Figure 5, E-G). Furthermore, we used qRT-PCR to further validate the expression levels of *HSP90-2* and *HSP17.4* in WT and *Zmheat1* mutants under control and heat-stress (45°C) 24h. Under control conditions, the expression of both genes did not differ significantly between WT and mutants. However, following heat stress, their expression was significantly downregulated in the *Zmheat1* mutant compared to WT (Figure 5, F and G). To verify whether ZmHEAT1 directly binds to the HSP promoter, we performed chromatin immunoprecipitation followed by qPCR (ChIP-qPCR) using an anti-Flag antibody in our *ZmHEAT1*-OE seedlings. We systematically analyzed the ∼2-kb promoter regions of two key differentially expressed HSP genes (*HSP90-2* and *HSP17.4*) with primer pairs spaced at 500-bp intervals. The results demonstrate significant, specific enrichment of ZmHEAT1 at distinct locations within the promoters of both target genes compared to the IgG control (Supplementary Figure S24, A and B). These data provide direct in vivo evidence that ZmHEAT1 physically associates with the promoters of these HSP genes, confirming its role as a direct transcriptional regulator in the heat stress response pathway. These data suggest that ZmHEAT1 positively regulates heat tolerance by affecting the integrity of the nuclear envelope and modulating the expression of a set of HTGs, including *HSP* and *HSF* genes.

## Discussion

In this study, we successfully used GWAS to explore favorable natural alleles associated with maize heat tolerance using the decrease in pollen viability during the reproductive stage under heat stress as a phenotype. From a panel of 257 inbred lines, we determined that *ZmHEAT1* has two main haplotypes and that Hap^T^, which is a favorable haplotype associated with a lower decrease in pollen viability under heat stress (Figure 1). This association was further validated by enhanced thermotolerance in a Hap^T^-carrying F□ segregating population, providing independent genetic evidence for the role of ZmHEAT1 in heat adaptation (Supplementary Figure S5). This natural allele of *ZmHEAT1*, which encodes the LINC-associated plant nuclear envelope protein ZmKAKU41. Pollen viability was dramatically lower in previously reported *Zmkaku41* mutants^21^. Here, we did not detect a decrease in pollen viability in the *Zmheat1* mutants under normal temperature conditions, but a lack of *ZmHEAT1* function decreased maize seed setting rate and yield under high temperature conditions (Figure 1). Our data further showed that ZmHEAT1/ZmKAKU41 improves maize heat tolerance at the seedling and reproductive stages (Figure 2). Thus, the Hap^T^ and *ZmHEAT1*-OE lines could be useful for genetic improvement of varieties that are sensitive to high temperature.

Nuclear envelope proteins play many important roles in cells, including maintaining nuclear structure and function, regulating transcription, DNA replication, nuclear envelope assembly, and participating in protein translation^16–18^. Nevertheless, little is known about the role of the nuclear envelope during heat stress responses. In Arabidopsis, yeast 2-hybrid evidence suggests there could be an interaction between AtKAKU4 (a homolog of ZmHEAT1/ZmKAKU41 and ZmKAKU42) and AtCRWN1 and AtCRWN4 proteins; our study fully demonstrated the interactions between ZmHEAT1 and ZmKAKU42, as well as NCH1 and NCH2 proteins by using systematic in vivo and in vitro experiments. ZmHEAT1 interacts with ZmKAKU42, as well as ZmNCH1 (but not ZmNCH2) to form a nuclear envelope complex critical for maintaining nuclear envelope integrity under heat stress (Figure 6). This mirrors the known interaction between AtKAKU4 and AtCRWN proteins in Arabidopsis, suggesting evolutionary conservation of this structural module. Meanwhile, given the interaction between KAKU4 and nucleoporins (NUP82/NUP136) in Arabidopsis^18^, it will be important to determine if a similar complex exists in maize, where it might regulate nucleoplasm-cytoplasm exchange and contribute to heat stress adaptation. Excessive accumulation of KAKU4 and CRWN/NCH proteins leads to nuclear envelope deformation and collapse^16,21^; in contrast, heat stress triggers their dissociation from the envelope and redistribution into the nucleoplasm^13^. Notably, we present a direct ultrastructural evidence through transmission electron microscopy demonstrating nuclear envelope rupture in *Zmheat1* mutants under heat stress, unequivocally establishing *ZmHEAT1* critical role in preserving nuclear envelope integrity during thermal challenge (Figure 5).

**Fig 6.**
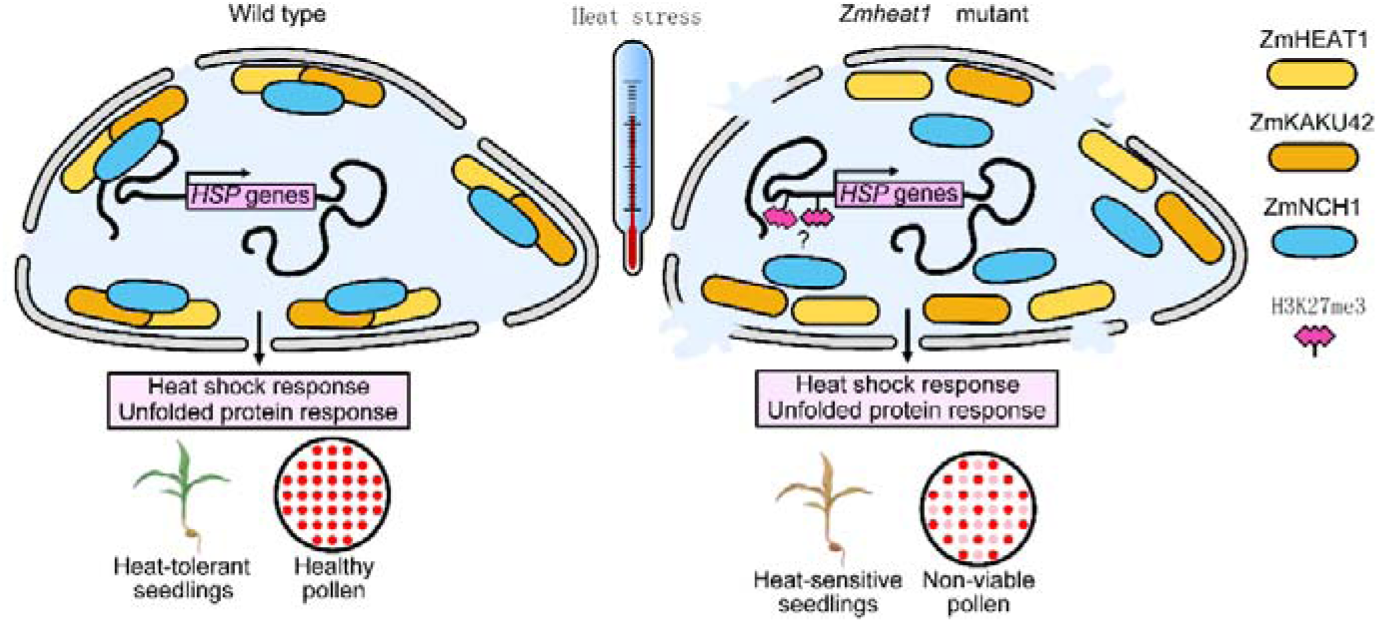
Proposed model of heat stress regulation by the ZmHEAT1/ZmKAKU42–ZmNCH1 complex in maize. In the wild type, ZmHEAT1, ZmKAKU42, and ZmNCH1 form a protein complex to maintain nuclear integrity and the shape of the nuclear envelope. In the *Zmheat1* or *Zmkaku42* mutant, the interaction of ZmHEAT1 or ZmKAKU42 with ZmNCH1 is disrupted, leading to the dissociation of ZmNCH1 from the nuclear periphery, resulting in a change in the shape of the nuclear envelope and damage to the nuclear envelope. The loss of ZmHEAT1 or ZmKAKU42 function also affects the expression of downstream *HSP*s, improving the heat tolerance of maize at the seedling and reproductive stages.

We acknowledge that the association signals detected at several loci, including *ZmHEAT1*, are of moderate strength. This likely reflects the inherent biological complexity of PVDR, a trait highly sensitive to environmental variation. Even subtle environmental fluctuations can markedly influence PVDR, thereby weakening detectable genetic effects and limiting the resolution of GWAS analyses. Such constraints are common in the genetic dissection of traits characterized by strong genotype-by-environment interactions. Regarding the effects of *ZmHEAT1* on the nuclear envelope under heat stress, initially, we attempted to examine mature pollen grains using TEM to visualize the nuclear envelope structure. However, the dense cytoplasmic contents obscured ultrastructural details, particularly of the nucleus and nuclear envelope, preventing clear visualization and reliable morphological assessment. Due to this technical limitation in pollen—and because *ZmHEAT1* and *ZmKAKU42* are also highly expressed in leaves and confer similar heat tolerance—we therefore utilized leaf tissue to assess nuclear envelope morphology. In addition, another two unresolved questions warrant further exploration: (1) Whether Hap^T^ and Hap^A^ alleles variations in *ZmHEAT1* induce post-translational modifications that alter protein functionality, and (2) The potential of *ZmHEAT1* alleles for agricultural applications. Crucially, field-based validation is required to determine whether Hap^T^ carriers and ZmHEAT1-overexpressing lines can maintain or enhance maize yield under realistic high-temperature conditions.

Under the general trend of global warming, it is crucial to explore heat-resistant genes related to the reproductive development stage for maize production. Not only should we pay attention to pollen, but also need to explore the development of silk and female ear. Moreover, since high temperature is often accompanied by drought and other biological stresses, it is imperative to carry out research related to compound stress. Overall, our work reveals a potential favorable haplotype for heat stress tolerance breeding in maize, and identifies ZmHEAT1, ZmKAKU42 and ZmNCH1 as important heat-stress–related factors, opening the door to understanding the role of the nuclear envelope in heat stress responses.

## Materials and Methods

### Plant growth conditions

All transgenic maize (*Zea mays*) plants and CRISPR/Cas9 mutants were generated by WIMI Biotechnology Co. Ltd. The constructs were transformed into the inbred line ‘B73-329’. Maize seeds were sown in pots (40 cm × 30 cm × 15 cm, length × width × depth) containing vermiculite and Pindstrup soil mix in a 1:1 (v:v) ratio; the plants were grown under a 16-h-light/8-h-dark photoperiod with 150 µmol m^−2^ s^−1^ white light with a 28□/22□ day/night temperature cycle and 40% relative humidity. Seeds of *Nicotiana benthamiana* were germinated in soil (1:1, vermiculite: Pindstrup), and seedlings were grown in the same growth chamber as above. Four-week-old *N. benthamiana* plants were used for experiments.

For heat treatment, T_2_ generation maize seedlings were grown in pots (40 cm × 30 cm × 15 cm, length × width × depth) to the V3 stage with water before being exposed to 45□ under a 16-h-light/8-h-dark photoperiod for 3–5 days using a heat chamber. The control seedlings were grown in a similar chamber but remained under control conditions. After heat treatment, the seedlings were allowed to recover under a 16-h-light/8-h-dark photoperiod with a 28□/22□ day/night temperature cycle for 3 days prior to taking photographs.

For Arabidopsis materials, seeds were stratified on 1/2 strength Murashige and Skoog (MS) agar medium containing 1.5% sucrose at 4°C for 2 days and then transferred to a growth chamber set at 22°C under a 16-hour light/8-hour dark photoperiod (light intensity, 110 μmol m^−2^ s^−1^, supplied by both incandescent and fluorescent lights) for 7 days. All transgenic lines were in the *Arabidopsis thaliana* Columbia ecotype background. Seeds from T_2_ generation overexpression-positive plants were harvested for heat treatment. 7-day old Arabidopsis seedlings were treated at 43°C heat stress for 1.5 hours, and the survival rate was calculated 7 days after recovery under normal temperature conditions.

### Phenotyping of heat tolerance and genome-wide association study (GWAS)

The 257 inbred lines of the maize population used in this study were planted at the Xiangshan Breeding Base of the Institute of Botany, Chinese Academy of Sciences, Beijing, in 2021. Pollen viability was determined during the anthesis stage. Specifically, tassels were manually cut in the field every morning, and pollen grains were collected by placing 30 μL of pollen grains into a 2-mL centrifuge tube, with another 30 μL of pollen grains transferred into a culture dish for incubation in a 41°C oven for 10 min. The AmphaZ32 pollen mass analyzer (Shanghai Zequan Technology Co., Ltd.) was used to determine pollen viability. The pollen viability decline rate was calculated using the following formula: pollen viability decline rate = ([pollen viability before treatment − pollen viability after treatment]/pollen viability before treatment) × 100%.

The GWAS was performed with the EMMAX algorithm using 558,659 high-quality SNPs (minor allele frequency ≥ 0.05) covering the entire maize genome^24–26^. The final significance threshold was set to *p* < 1.03 × 10^−5^ (1/97,438) for the GWAS based on 97,438 independent SNPs by filtering SNPs in strong LD using PLINK (window size 50, step size 50, r^2^ ≥ 0.2).

### Physiological assays

To detect H_2_O_2_ accumulation in leaves, 3,3′-diaminobenzidine (DAB) staining was used as described in a previous study^27^. Briefly, V3-stage maize seedlings grown under control conditions were exposed to 45□ for 24 h before being transferred to a 1 mg/mL DAB solution and vacuum-infiltrated for 30 min before incubation for 10 h at room temperature. Subsequently, the leaves were cleared in 95% (v/v) ethanol until all chlorophylls were leached out of the samples.

### Plasmid construction and plant transformation

The transgenic lines used in this study were generated in the maize inbred line B73-329, which was selected as the recipient material due to its widespread use and high amenability to genetic transformation. The transformation constructs were generated by cloning the full-length *ZmHEAT1* coding sequence into the pWMV013 vector (derived from pCAMBIA3300, obtained from WIMI Biotechnology Co. Ltd.). To obtain mutants of *ZmHEAT1* and *ZmKAKU42* using CRISPR/Cas9-mediated genome editing, fragments of the first exon were selected as target sites of single guide RNAs (http://cbi.hzau.edu.cn/cgi-bin/CRISPR), whose sequences were cloned into the pCPB-ZmUbi-Cas9 vector at the HindIII restriction site^28^. Transgenic plants were obtained by Agrobacterium (*Agrobacterium tumefaciens*)-mediated transformation as previously described^29^.

To generate the *ZmNCH1*-OE line in Arabidopsis, the full-length of ZmNCH1 coding sequence were cloned into pEarleyGate 301 vector. Transformation of *Arabidopsis thaliana* was performed by the floral dip method^30^. Transgenic plants were isolated on 1/2 MS medium containing Basta (10 μg/ml), and positive seedlings were transferred to soil. The seeds from the T_2_ generation of positive transgenic plants were used for further analysis.

### RT-qPCR analysis of gene expression in different transgenic maize lines

Total RNA was extracted from the leaves of V3-stage maize seedlings using a plant RNA kit (ER301; TransGen Biotech, China). Total RNA (1 μg) was reverse transcribed into first-strand complementary DNA (cDNA) using M-MLV reverse transcriptase according to the manufacturer’s instructions (Vazyme, China). qPCR analysis was performed as previously described^31^. *ACTIN* was used as internal reference to normalize the expression value of each sample using the ΔΔCt method. Primers used for RT-qPCR are listed in Supplementary Data Set S5. The experiments were performed independently with three technical replicates for each of three independent biological replicates from different seedlings.

### ChIP-qPCR

The ChIP assay was performed as previously described with minor modifications^32^. *ZmNCH1*-OE plants of fourteen-day-old after sowing were harvested in the chamber and fixed with 1% formaldehyde under a vacuum for 10 min. The reaction was quenched by additional glycine to a final concentration of 0.125 M, and the mixture was incubated for another 5 min under vacuum at room temperature. The chromatin was isolated and sonicated, and DNA fragments associated with ZmNCH1-Flag proteins were co-immunoprecipitated using antibody of anti-Flag or anti-IgG, respectively. Each treatment had two biological replicates. The enrichment of DNA fragments was quantified by qPCR. Related qPCR primers were listed in the Supplementary Data Set S5.

### Yeast two-hybrid assays

The full-length coding sequences of *ZmHEAT1* and *ZmKAKU42* were cloned individually into the pGBKT7 vector (Clontech) to generate the BD-*ZmHEAT1* and BD-*ZmKAKU42* constructs. The full-length coding sequences of *ZmKAKU42*, *ZmNCH1*, and *ZmNCH2* were cloned individually into the pGADT7 vector (Clontech) to generate the AD-*ZmKAKU42*, AD-*ZmNCH1*, and AD-*ZmNCH2* constructs, respectively. The resulting constructs were co-transformed as appropriate pairs into the yeast (*Saccharomyces cerevisiae*) strain Y2HGold; the transformed cells were plated on synthetic defined (SD) medium lacking leucine and tryptophan (−Trp−Leu) and grown at 30°C for 3 days. The positive transformants were then evaluated for growth on SD medium lacking Leu, Trp, histidine, and adenine (SD/−Trp−Leu−His−Ade) at 30°C for 5 days. Empty pGADT7 and pGBKT7 vectors were co-transformed as a negative control.

### Bimolecular fluorescence complementation assays

The vectors pSPYNE-35S and pSPYCE-35S encoding the N-terminal half of YFP (nYFP) or the C-terminal half of YFP (cYFP), respectively, were used for co-infiltration of *N. benthamiana* leaves via Agrobacterium^33^. The full-length *ZmHEAT1* and *ZmKAKU42* coding sequences were cloned individually into pSPYCE-35S. The full-length *ZmKAKU42*, *ZmNCH1*, and *ZmNCH2* coding sequences were cloned individually in pSPYNE-35S. The resulting constructs were co-infiltrated as appropriate pairs into *N. benthamiana* leaves, which were then incubated in chambers for 48 h to allow transgene expression. The YFP fluorescence signal was detected under a Zeiss LSM 980 confocal microscope with an excitation wavelength of 488 mm and an emission wavelength of 518–582 nm.

### GST pull-down assays

The full-length coding sequences of *ZmHEAT1* and *ZmKAKU42* were cloned individually into pGEX□4T□1 to obtain the recombinant proteins ZmHEAT1□GST and ZmKAKU42□GST, respectively, while the full-length coding sequence of *ZmKAKU42* was cloned into pMAL-c5x to obtain recombinant ZmKAKU42□MBP. The full-length coding sequence of *ZmNCH1* was cloned into pET32a to obtain recombinant ZmNCH1-His. The recombinant proteins ZmHEAT1□GST, ZmKAKU42□GST, and GST were produced in *Escherichia coli* Rosetta 2 (DE3) by the addition of 0.2 mM IPTG and incubation at 16°C for 20 h; the cells were collected and broken with an JY96-IIN Ultrasonic Homogenizer (Ningbo Scientz Biotechnology Co., Ltd) and purified using glutathione beads (BeaverBeads). ZmNCH1-His was produced in Rosetta 2 (DE3) by the addition of 0.2 mM IPTG and incubation at 16°C for 24 h and purified using IDA-Nickel (BeaverBeads). ZmKAKU42□MBP was produced in Rosetta 2 (DE3) by the addition of 0.2 mM IPTG, incubated at 16°C for 24 h, and purified using 6FF dextrin beads (LABLEAD). Recombinant purified ZmHEAT1□GST, ZmKAKU42□GST, or GST was incubated with glutathione beads at 4°C for 2.5 h. Recombinant purified ZmKAKU42□MBP or ZmNCH1-His was then added to the mixture, followed by incubation for 1.5 h at 4°C. The glutathione beads and bound proteins were washed six times with phosphate-buffered saline (PBS) for 5 min each time. The proteins bound to the beads were subjected to SDS-PAGE analysis followed by immunoblot analysis with anti-MBP (1:5,000 dilution; ABclonal) and anti-His antibodies (1:5,000 dilution; ABclonal).

### Coimmunoprecipitation assays

The full-length coding sequences of *ZmHEAT1* and *ZmKAKU42* were cloned individually into the pCAMBIA1300-GFP vector at the PstI and KpnI restriction sites, while the full-length coding sequences of *ZmHEAT1* and *ZmNCH1* were cloned individually into the pCAMBIA1300-Flag vector at the HindIII and SpeI restriction sites^34^. The resulting constructs were transformed into cells of Agrobacterium strain GV3101. Cultures of positive colonies were resuspended in infiltration buffer (10 mM MES-KOH, 10 mM MgCl_2_, and 0.2 mM acetosyringone) and mixed in a 1:1 (v/v) ratio before being co-infiltrated into the leaves of 4-week-old *N. benthamiana* plants. After a 48-h incubation, leaf samples were collected, ground into fine powder in liquid nitrogen, and homogenized in extraction buffer (50 mM Tris-HCl pH 7.5, 150 mM NaCl, 1 mM EDTA, 1% [v/v] Triton X-100, 10% [v/v] glycerol, 1×protease inhibitor cocktail [Roche, #04693159001], and 50 µM MG132). Immunoprecipitation was performed using Magnetic Beads-conjugated mouse anti-GFP-Tag mAb (ABclonal) at 4°C for 2 h and then the beads were washed six times with washing buffer (50 mM Tris-HCl pH 7.5, 150 mM NaCl, 1 mM EDTA, 1% [v/v] Triton X-100, and 10% [v/v] glycerol). The immunoprecipitates were separated by SDS-PAGE and detected by immunoblot with anti-GFP (1:5,000 dilution; ABclonal) or anti-Flag (1:5,000 dilution; ABclonal) antibodies.

### Subcellular localization

The ZmHEAT1-GFP, ZmKAKU42-GFP, ZmNCH1-GFP, and ZmNCH2-GFP constructs generated above in the pCAMBIA1300-GFP vector were individually introduced into Agrobacterium strain GV3101. Bacterial cultures from positive colonies were resuspended in infiltration buffer (10 mM MgCl_2_, 10 mM MES pH 5.6, and 150 µM acetosyringone) to an OD at 600 nm of 0.8 and then infiltrated into the leaves of *N. benthamiana* plants via Agrobacterium-mediated infiltration. The infiltrated plants were cultured at 22°C for 48 h before being exposed to heat stress at 45°C for 1 h. GFP signals were collected under a Zeiss LSM 980 confocal microscope with an excitation wavelength of 488 nm and an emission wavelength of 513 nm. Agrobacterium containing H2B-mCherry was used as a nuclear localization marker^35^.

### Determination of mature pollen viability

To explore the association analysis of *ZmHEAT1* with the pollen viability decline rate in maize, mature pollen of *Zmheat1* knockout lines was treated at room temperature or 41□ for 30 min in an oven. Pollen viability was evaluated by 2,3,5-triphenyl tetrazolium chloride (TTC) staining. The TTC solution (8 g/L) was prepared in 1 M potassium phosphate buffer, pH 7.0. The staining of pollen grains was then observed using a SZ680 series zoom stereomicroscope (CHONGQING OPTEC INSTRUMENT CO., LTD), and at least three fields of view were captured for each sample. Pollen viability was scored as the ratio of stained pollen grains to total pollen grains.

### RNA-seq analysis

For RNA-seq, the leaves and stems of V3-stage maize seedlings (*Zmkaku41-1* and WT) exposed to heat at 45°C for 24 h were collected for extraction of total RNA with TRIzol reagent (Invitrogen, USA). The sequencing libraries were constructed using 1 μg total RNA per sample. Libraries were prepared using a TruSeq RNA Library Preparation kit (Illumina, USA) and sequenced on an Illumina Novaseq platform, as 150-bp paired-end reads. fastp^36^ was used to remove the adaptor sequences, low-quality bases (Q30), and sequences containing >10% undetermined bases. The resulting clean reads were aligned to the maize reference genome B73_AGPv4^37^ using Hisat2^38^ with default parameters. Subsequently, Stringtie^39^ was used to quantify the uniquely aligned reads (as FPKM), and the Pearson’s correlation coefficient method was used to evaluate the pairwise reproducibility between samples. Principal component analysis and identification of differentially expressed genes (DEGs) was carried out using the R package DESeq2 (v.1.30.0). *p*-value were adjusted using the Benjamini-Hochberg procedure. DEGs were selected based on the following criteria: *p* < 0.05 and log_2_ (fold-change) ≥ 1. Three independent replicates were performed for each sample at each time point. Cluster analysis was performed using the R package pheatmap (v.1.0.12; https://cran.r-project.org/web/packages/pheatmap/index.html). Functional enrichment categories in clusters were analyzed using GoSlim with all genes as background. Based on results from a hypergeometric test, GO terms with false discovery rate values < 0.05 were considered significantly enriched.

### Transmission electron microscopy

The ultrastructure of maize leaf cell nuclear was determined by transmission electron microscopy. The leaf tissues were cut into 1.0 × 3.0 mm pieces and fixed in 3% (w/v) glutaraldehyde in 0.1 M PBS, pH 7.2, at 4°C overnight. The fixed tissues were washed in PBS three times for 20 min each at 4°C, post-fixed for 4 h in 1% (w/v) osmium tetroxide, dehydrated in a graded acetone series, and infiltrated with Spurr resin (SPI Supplies USA), which was allowed to polymerize at 60°C for 24 h. The samples were cut into ultrathin sections (80-nm thickness) with a Leica Ultracut R (Germany) and examined using a transmission electron microscope (H-7700; Hitachi, Japan) at 80 kV.

### Statistical methods

Differences between two groups were assessed using a two-sided Student’s *t*-test. For multiple comparisons, significance analysis was performed with one-way analysis of variance (ANOVA) followed by Tukey’s multiple comparison tests. All statistical analyses were performed using GraphPad Prism version 8.08 and are summarized in Supplementary Data Set S6.

## Supporting information

Genes differentially expressed in the WT plants with and without heat treatment, related to Figure 5C

Genes differentially expressed under normal conditions, Zmheat1 versus WT, related to Figure S22 B and C

Genes differentially expressed with heat treatment, Zmheat1 versus WT, related to Figure S22 B and C

TPM value of 1,812 ZmHEAT1-regulated HTG genes, related to Figure S23

Statistical analyses

## Accession numbers

Genes mentioned in the study are as follows: *ZmHEAT1* (Zm00001d026487), *ZmKAKU42* (Zm00001d002012), *ZmNCH1* (Zm00001d051600), *ZmNCH2* (Zm00001d043335), *ZmHSP17.4* (Zm00001d039936), *ZmHSP90-2* (Zm00001d031332), and *ZmACTIN* (Zm00001d010159).

## Acknowledgments

We gratefully thank Dr. Feng-Qin Dong (Institute of Botany) and Dr. Jing-Quan Li (Institute of Botany) for technical assistance, we also thank Prof. Mingliang Xu (China Agricultural University) provide F_2_ population.

## Author contributions

M.Z. conceived the project and designed the experiments. Y.L.J. and S.F.W. carried out most of the experiments; X.L.W. performed most data analyses; H.L., Z.L., W.C., Y.J.H and H.M.S. carried out some experiments. Y.L.J. and S.F.W. wrote the draft. X.HY., P.H.W. and M.Z. revised the manuscript. All authors have read and approved the final manuscript.

## Data availability

The data generated in this study has been uploaded to the NCBI database and can be retrieved under accession number PRJNA1233201. All scripts used for quantification, normalization and statistical testing, are available in the Github: https://github.com/WangXiaoLiang1108/RNASEQ-FOR-ZmHEAT1.

## Supplemental information

Document S1. Supplementary Figure. S1-S24, Supplemental Data Set S1-S6, and supplemental references

