## Supplemental Data 1 for "The ZmHEAT1/ZmKAKU42–ZmNCH1 module regulates heat stress tolerance by modulating nuclear envelope structure in maize"

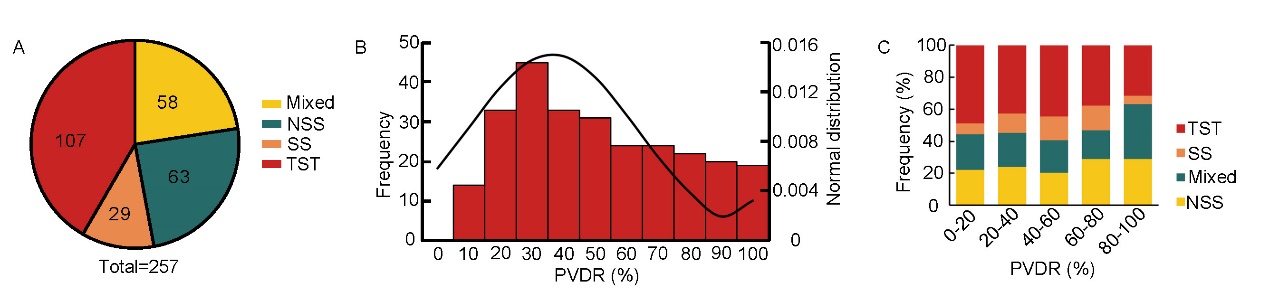

**Figure. S1** Distribution of pollen viability decline rate among maize inbred lines.

A. Distribution of the 257 inbred lines from germplasm collections across four subpopulations: TST (tropical/subtropical lines), NSS (non-stiff stalk), SS (stiff stalk), and Mixed. NSS and SS lines are temperate lines; SS lines are derived from B73; Mixed lines are of mixed origin.
B. Distribution of the maize pollen viability decline rate (PVDR) in a heat tolerance test of 257 inbred lines.
C. Frequency distribution of heat tolerance for lines from each of the four subpopulations, as revealed by their associated PVDR.

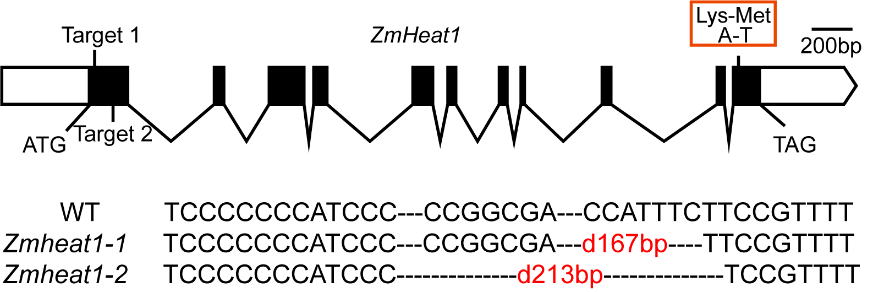

**Figure S2.** Construction of CRISPR/Cas9-mediated *Zmheat1* knockout lines. Two sgRNAs that specifically target *ZmHEAT1* were designed. Two mutant lines were identified, *Zmheat1-1* and *Zmheat1-2*. Black rectangles indicate exons, white rectangles indicate untranslated regions (UTRs), and black lines connecting rectangles indicate introns. A natural allele with an A-to-T substitution in exon 11 results in a lysine-to-methionine change.

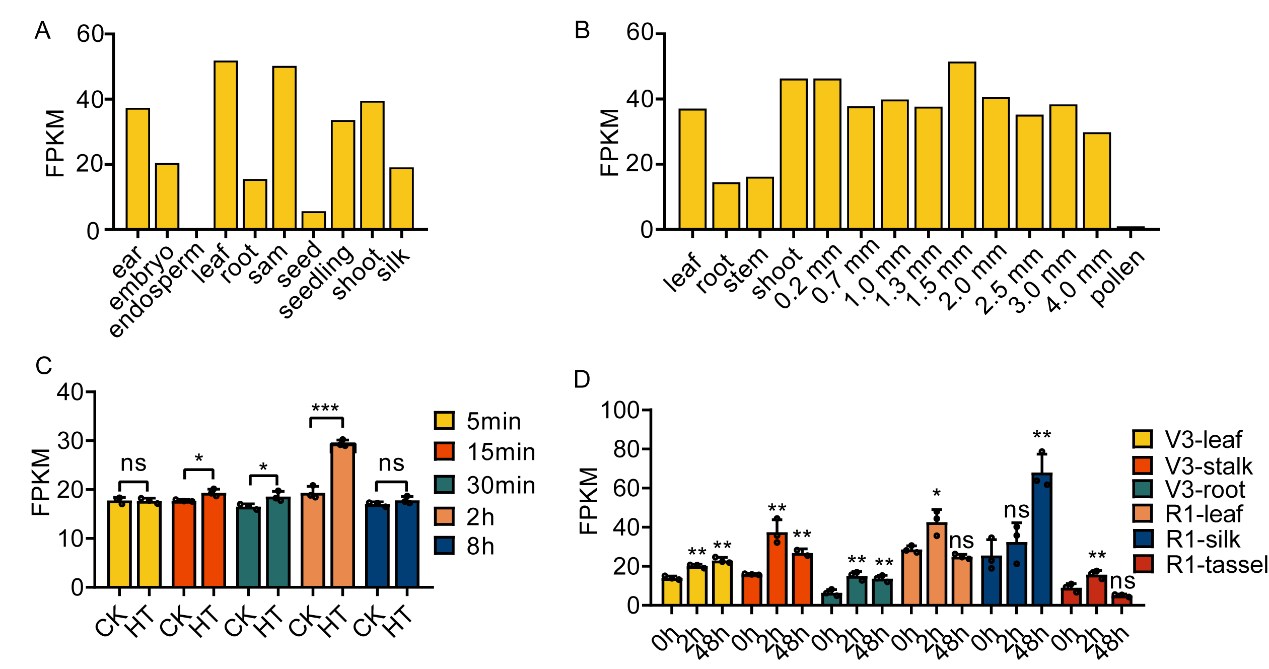
 **Figure S3.** Analysis of *ZmHEAT1* expression pattern.

A. Relative *ZmHEAT1* expression levels in different tissues of the maize inbred line B73^1-4^.
B. Relative *ZmHEAT1* expression levels in different tissues of the maize inbred line Chang7-2. Samples 0.2 mm, 0.7 mm, 1.0 mm, 1.3 mm, 1.5 mm, 2.0 mm, 2.5 mm, 3.0 mm, and 4.0 mm represent anthers of different lengths^5^.
C. Relative *ZmHEAT1* expression levels in the maize inbred line B73 after high temperature treatment at 45℃ for 5 min, 15 min, 30 min, 2 h, or 8 h^6^. CK, control; HT, heat treatment. Statistical significance was determined by a two-sided *t*-test; *, *p* < 0.05; ***, *p* < 0.001.
D. Expression of *ZmHEAT1* in leaf, stem, and root tissues at the three-leaf stage and in ear, tassel, leaf, and silk tissues at the silking stage under high temperature treatment^6^. Statistical significance was determined by one-way ANOVA; *, *p* < 0.05; **, *p* < 0.01. For (C, D), values are means ± SD with all individual data points shown as black dots.

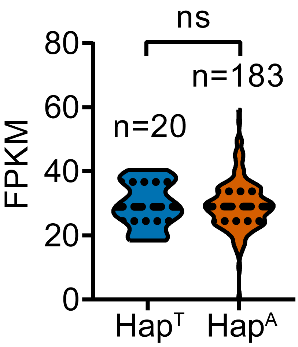

**Figure S4.** Expression levels of *ZmHEAT1* in the V3-seedling leaves of Hap^T^ and Hap^A^. Statistical significance was determined by a two-sided *t*-test (n=20 for Hap^T^, n=183 for Hap^A^; ns, not significant)^7^.

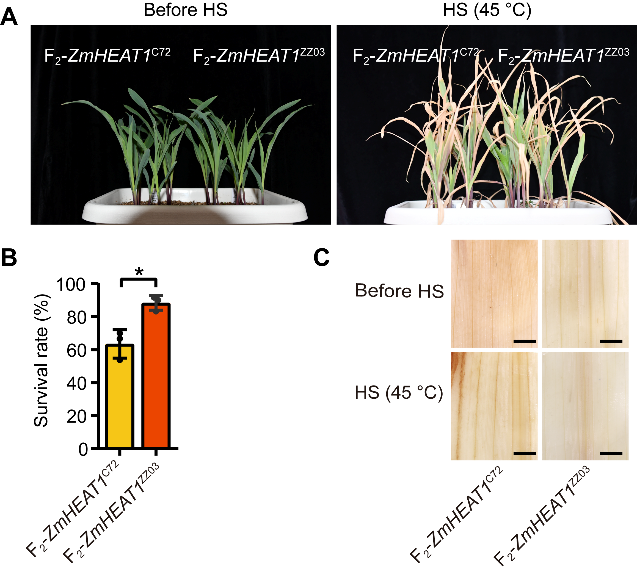

**Figure. S5** Identification of seedling thermotolerance in natural variation.

A. Representative photographs of V3-stage seedlings of the F_2_-*ZmHEAT1*^C72^ and F_2_-*ZmHEAT1*^ZZ03^ grown under control conditions (left) and after treatment at 45℃ for 5 days and recovery at 28℃/22℃ for 3 days (right).
B. Survival rate of F_2_-*ZmHEAT1*^C72^ and F_2_-*ZmHEAT1*^ZZ03^ seedlings after recovery from HS at 28℃/22℃ for 3 days. Statistical significance was determined by one-way ANOVA; *, *p* < 0.05. Values are means ± SD with all individual data points shown as black dots.
C. DAB staining of leaves from F_2_-*ZmHEAT1*^C72^ and F_2_-*ZmHEAT1*^ZZ03^ V3-stage seedlings grown at 28℃/22℃ (Before HS, top), or exposed to 45℃ for 24 h (bottom). Scale bars, 100 µm.

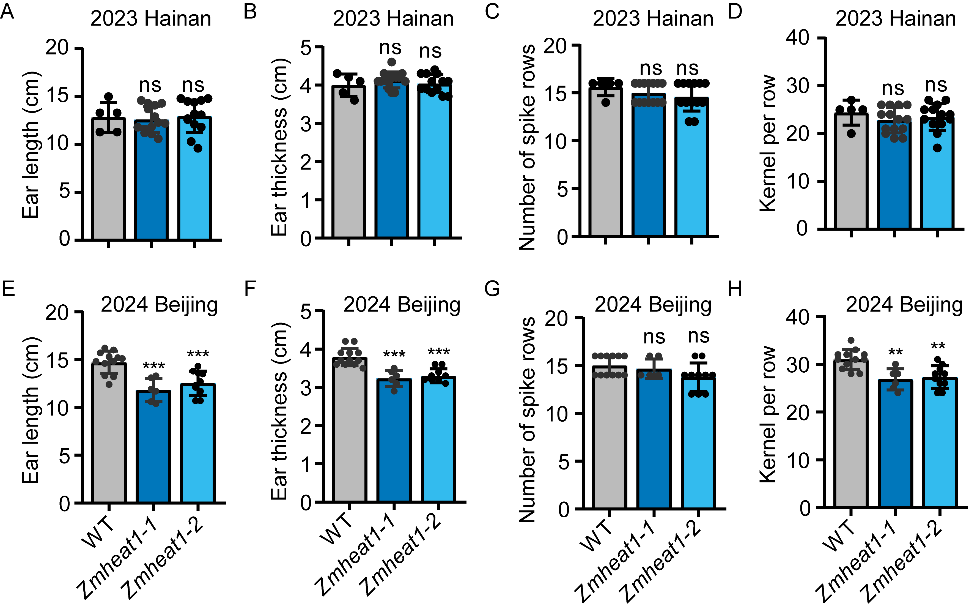

**Figure S6.** Effects of high temperature on ear traits of the wild type and *Zmheat1* mutants.

A–H. Ear length (A, E), ear thickness (B, F), number of spike rows (C, G), and kernels per row (D, H) of the wild type and *Zmheat1-1* and *Zmheat1-2* grown under control conditions in Hainan in 2023 (A–D) or under heat stress Beijing in 2024 (E–H). Statistical significance was determined by one-way ANOVA; *, *p* < 0.05; **, *p* < 0.01; ***, *p* < 0.001. Values are means ± SD with all individual data points shown as black dots.

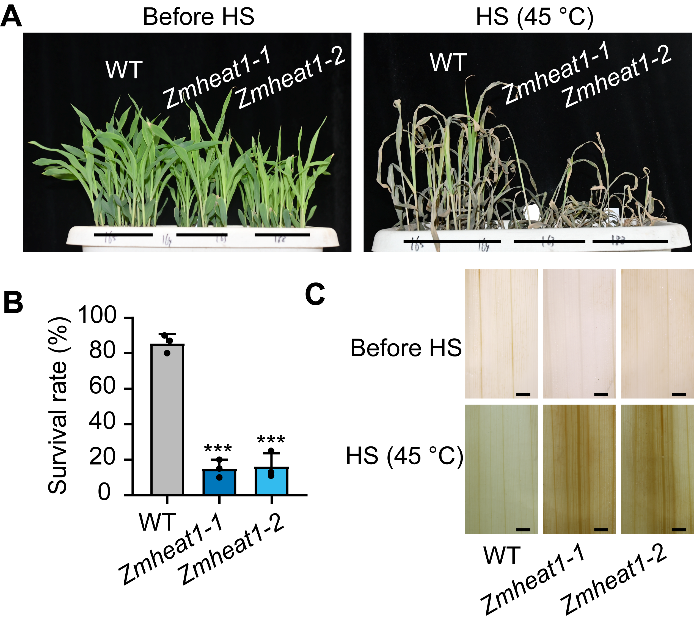

**Figure S7.** Effects of high temperature on seedling of the wild type and *Zmheat1* mutants.

A. Representative photographs of V3-stage seedlings of the wild type and *Zmheat1* mutants grown under control conditions (left) and after treatment at 45℃ for 3 days and recovery at 28℃/22℃ for 3 days (right).
B. Survival rate of wild-type and *Zmheat1* mutant seedlings after recovery from HS at 28℃/22℃ for 3 days. Statistical significance was determined by one-way ANOVA; ***, *p* < 0.001. Values are means ± SD with all individual data points shown as black dots.
C. DAB staining of leaves from wild-type and *Zmheat1* V3-stage seedlings grown at 28℃/22℃ (Before HS, top), or exposed to 45℃ for 24 h (bottom). Scale bars, 100 µm.

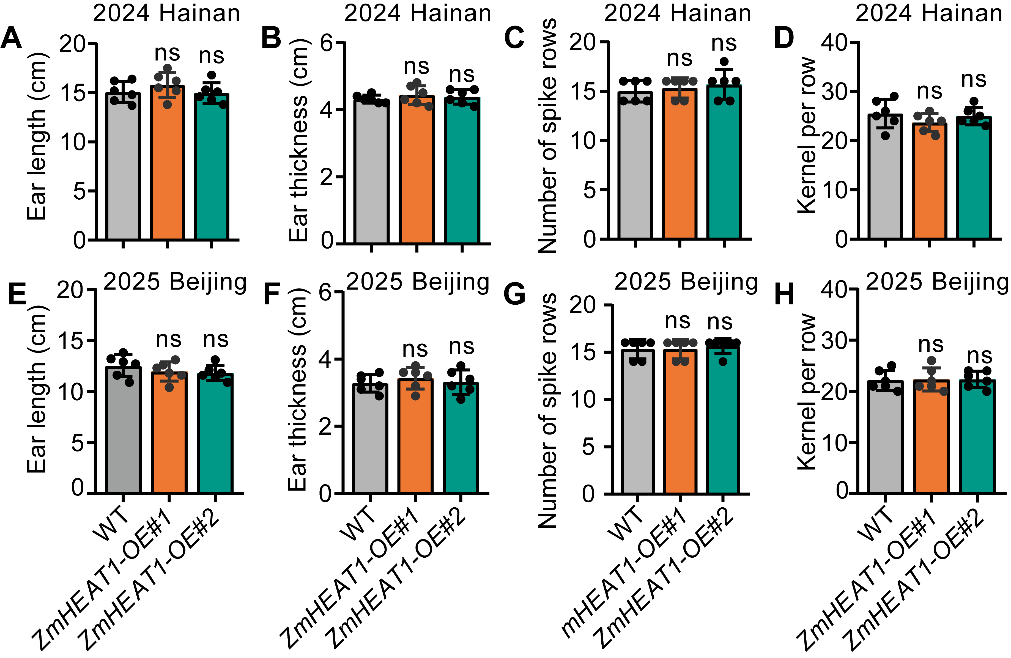

**Figure S8.** Effects of high temperature on ear traits of the wild type and *ZmHEAT1*-OE mutants.

A–H. Ear length (A, E), ear thickness (B, F), number of spike rows (C, G), and kernels per row (D, H) of the wild type and *ZmHEAT1*-OE#1 and *ZmHEAT1*-OE#2 grown under control conditions in Hainan in 2024 (A–D) or under heat stress Beijing in 2025 (E–H). Statistical significance was determined by one-way ANOVA; ns, not significant. Values are means ± SD with all individual data points shown as black dots.

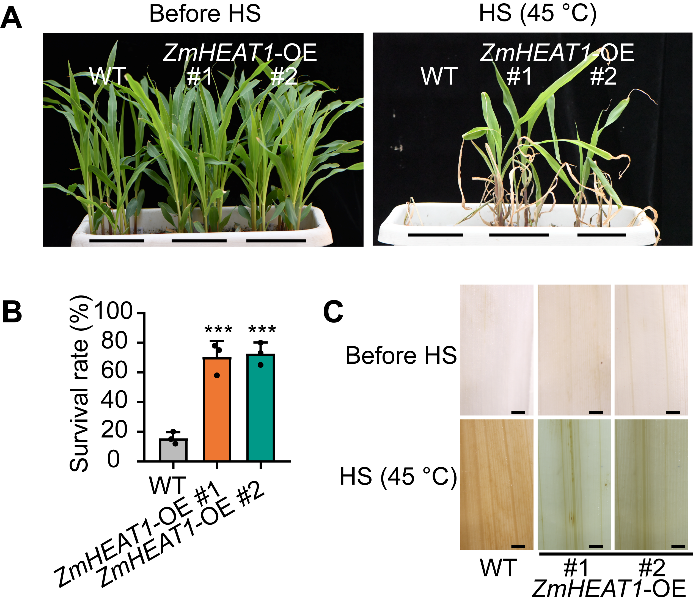

**Figure S9.** Effects of high temperature on seedling of the wild type and *ZmHEAT1*-OE mutants.

A. Representative photographs of V3-stage seedlings of the wild type and *ZmHEAT1*-OE mutants grown under control conditions (left) and after treatment at 45℃ for 5 days and recovery at 28℃/22℃ for 3 days (right).
B. Survival rate of wild-type and *ZmHEAT1*-OE mutant seedlings after recovery from HS at 28℃/22℃ for 3 days. Statistical significance was determined by one-way ANOVA; ***, *p* < 0.001. Values are means ± SD with all individual data points shown as black dots.
C. DAB staining of leaves from wild-type and *ZmHEAT1*-OE V3-stage seedlings grown at 28℃/22℃ (Before HS, top), or exposed to 45℃ for 24 h (bottom). Scale bars, 100 µm.

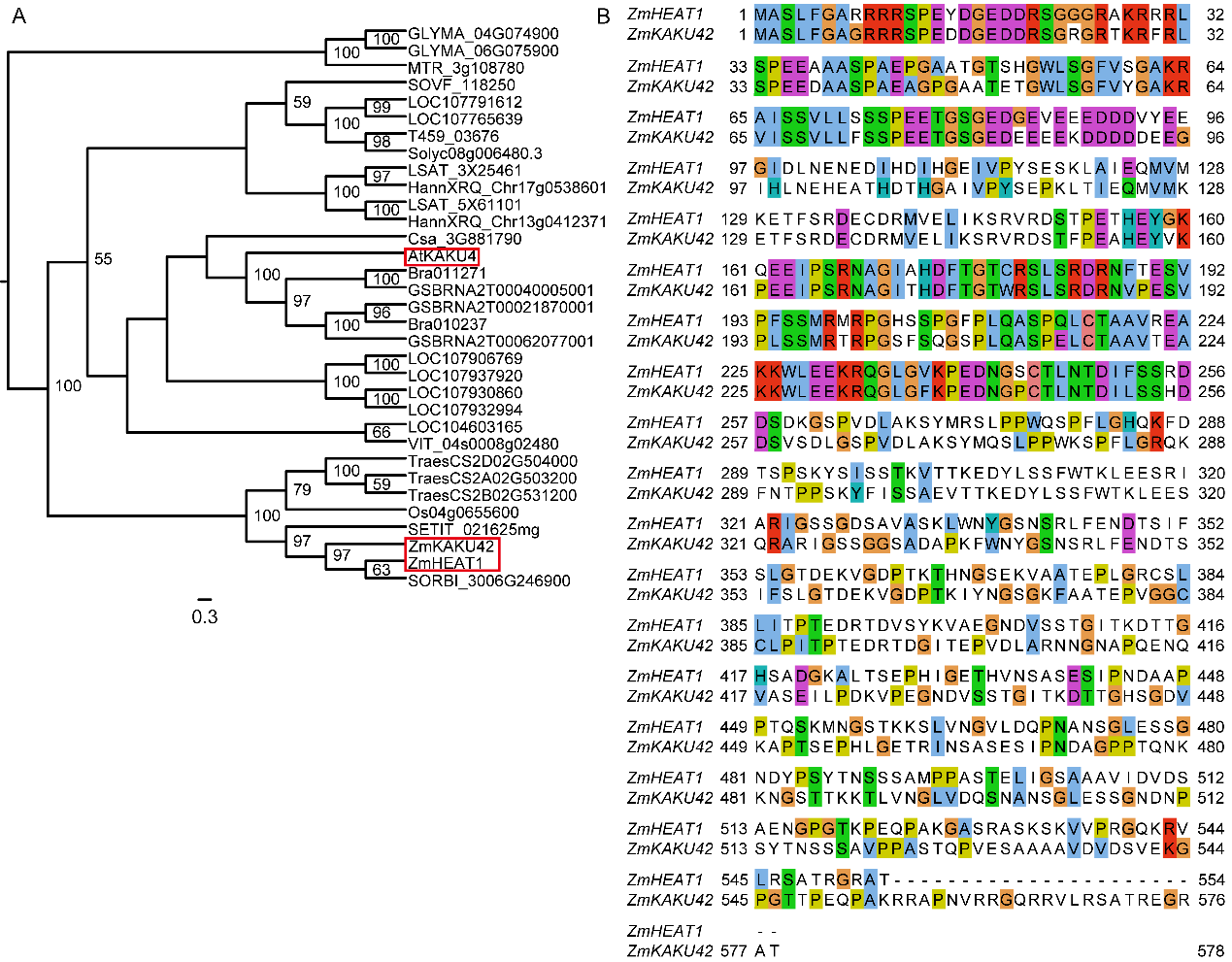

**Figure S10.** ZmHEAT1 and ZmKAKU42 are highly similar.

A. Phylogenetic tree showing the evolutionary relationship of *ZmHEAT1* (encoded by *Zm00001d026487*) and its homologs in various plant species. The horizonal lines are branches and represent evolutionary lineages changing over time. The longer the branch in the horizonal dimension, the larger the amount of change. The bar at the bottom of the figure provides a scale for this.
B. Amino acid sequence alignment of *ZmHEAT1* and *ZmKAKU42* (encoded by *Zm00001d002012*).

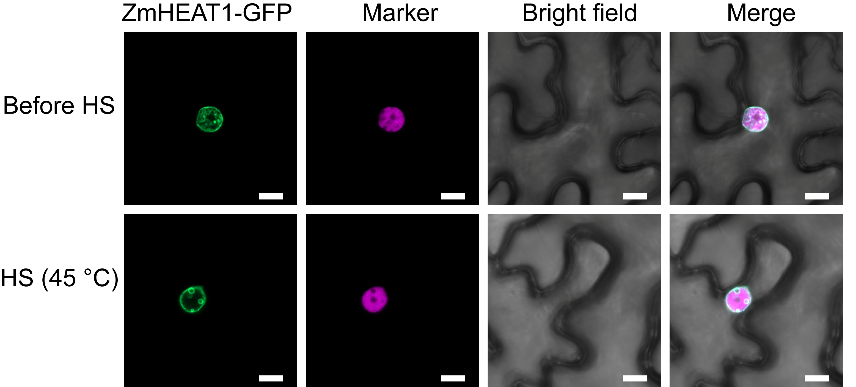

**Figure S11.** Subcellular localization of ZmHEAT1-GFP in *N. benthamiana* leaves. The nuclear marker was histone H2B-mCherry. Scale bars, 5 µm.

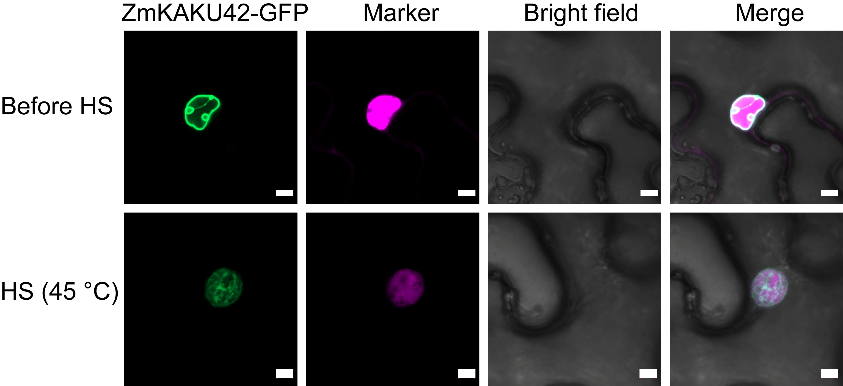

**Figure S12.** Subcellular localization of ZmKAKU42-GFP in *N. benthamiana* leaves. The nuclear marker was histone H2B-mCherry. Scale bars, 5 µm.

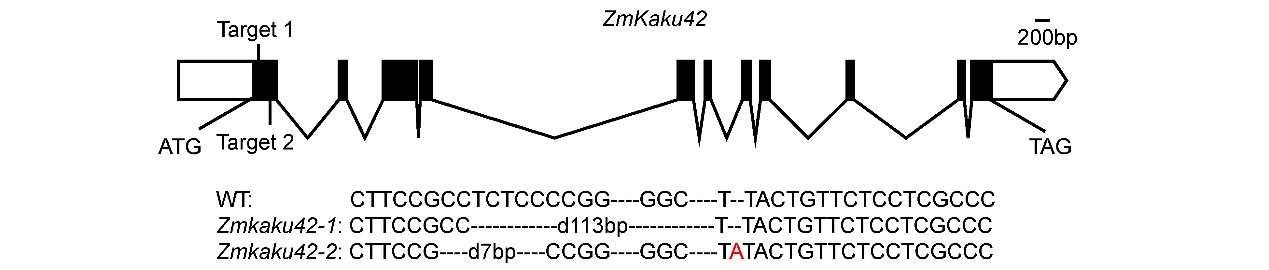

**Figure S13.** Construction of CRISPR/Cas9-mediated *Zmkaku42* knockout lines. Two sgRNAs that specifically target *ZmKAKU42* were designed. Two mutants were identified: *Zmkaku42-1* and *Zmkaku42-2*. Black rectangles indicate exons, white rectangles indicate UTRs, and black lines indicate introns.

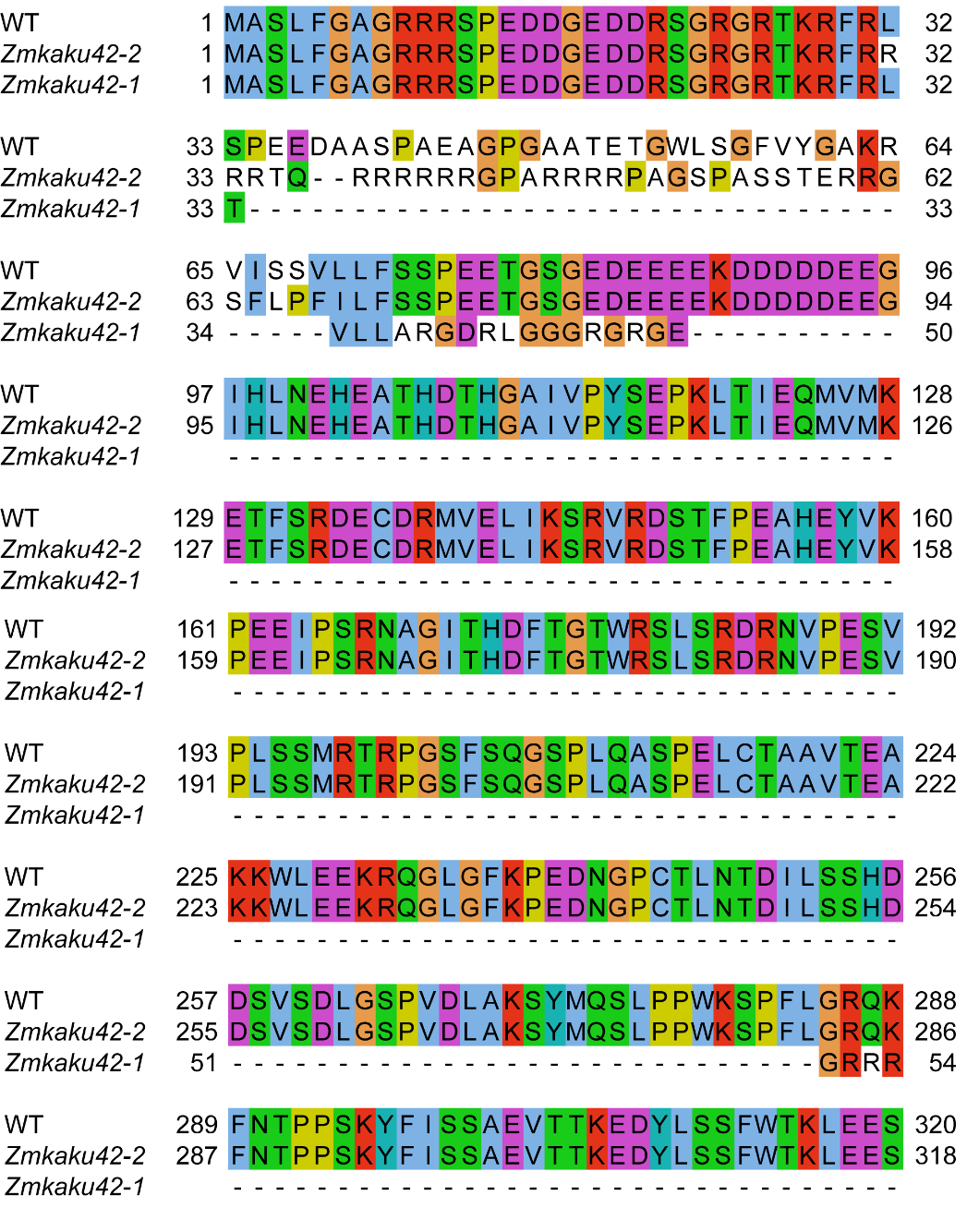

**Figure S14.** Comparison of predicted protein sequences of *Zmkaku42-1* and *Zmkaku42-2* with WT.

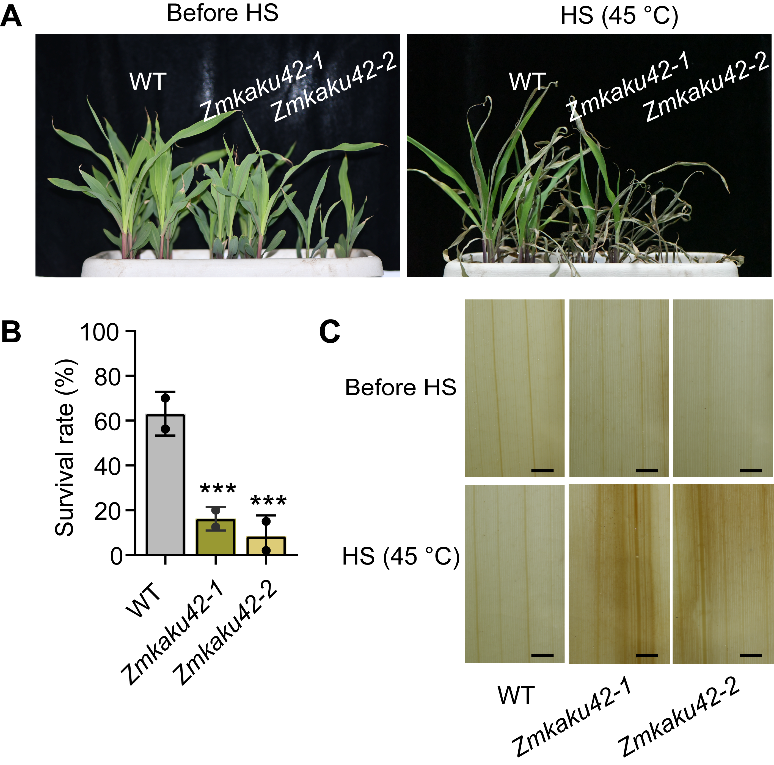

**Figure S15.** Effects of high temperature on seedling of the wild type and *Zmkaku42* mutants.

A. Representative photographs of V3-stage seedlings of the wild type and *Zmkaku42* mutants grown under control conditions (left) and after treatment at 45℃ for 3 days and recovery at 28℃/22℃ for 3 days (right).
B. Survival rate of wild-type and *Zmkaku42* mutant seedlings after recovery from HS at 28℃/22℃ for 3 days. Statistical significance was determined by one-way ANOVA; ***, *p* < 0.001. Values are means ± SD with all individual data points shown as black dots.
C. DAB staining of leaves from wild-type and *Zmkaku42* V3-stage seedlings grown at 28℃/22℃ (Before HS, top), or exposed to 45℃ for 24 h (bottom). Scale bars, 100 µm.

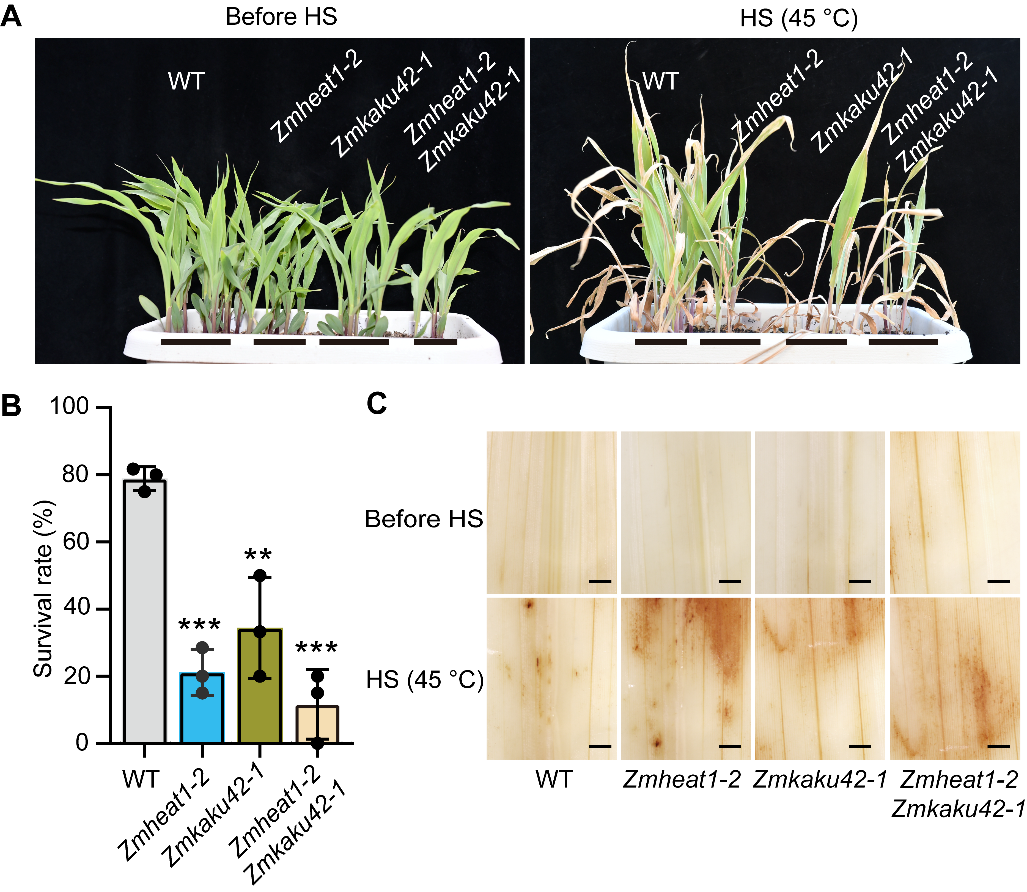

**Figure S16.** Effects of high temperature on seedling of the wild type, *Zmheat1*, *Zmkaku42* and *Zmheat1 Zmkaku42* mutants.

A. Representative photographs of V3-stage seedlings of the wild type wild type, *Zmheat1-2*, *Zmkaku42-1* and *Zmheat1-2 Zmkaku42-1* mutants grown under control conditions (left) and after treatment at 45℃ for 3 days and recovery at 28℃/22℃ for 3 days (right).
B. Survival rate of wild type, *Zmheat1-2*, *Zmkaku42-1* and *Zmheat1-2 Zmkaku42-1* mutant seedlings after recovery from HS at 28℃/22℃ for 3 days. Statistical significance was determined by one-way ANOVA; ***, *p* < 0.001, **, *p* < 0.01. Values are means ± SD with all individual data points shown as black dots.
C. DAB staining of leaves from wild type, *Zmheat1-2*, *Zmkaku42-1* and *Zmheat1-2* *Zmkaku42-1* V3-stage seedlings grown at 28℃/22℃ (Before HS, top), or exposed to 45℃ for 24 h (bottom). Scale bars, 100 µm.

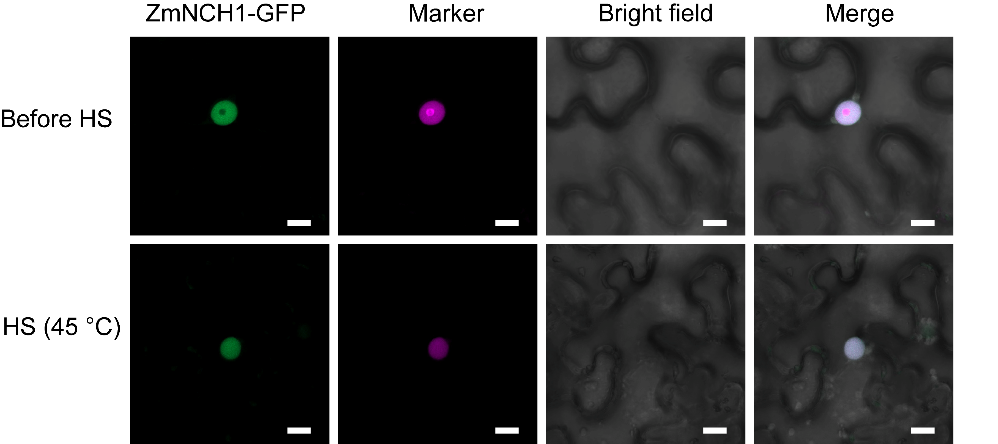

**Figure S17.** Subcellular localization of ZmNCH1-GFP in *N. benthamiana* leaves. The nuclear marker was histone H2B-mCherry. Scale bars, 5 µm.

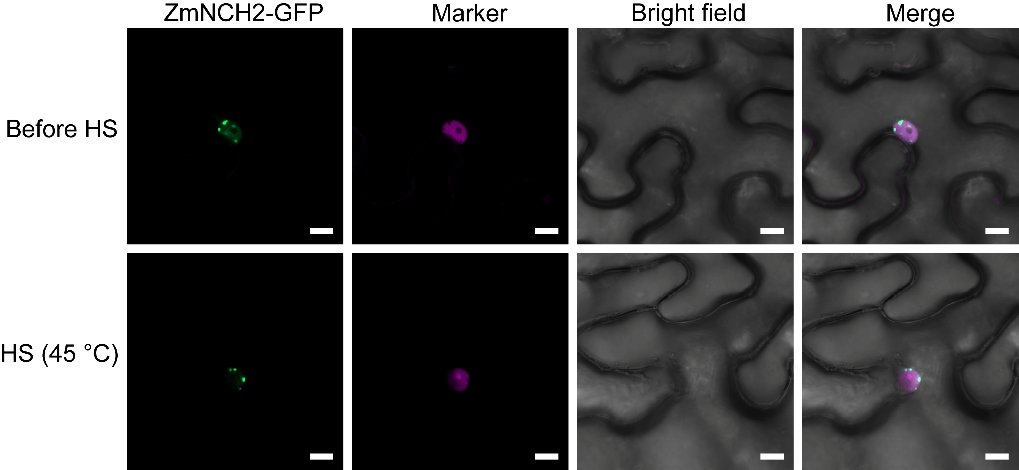

**Figure S18.** Subcellular localization of ZmNCH2-GFP in *N. benthamiana* leaves. The nuclear marker was histone H2B-mCherry. Scale bars, 5 µm.

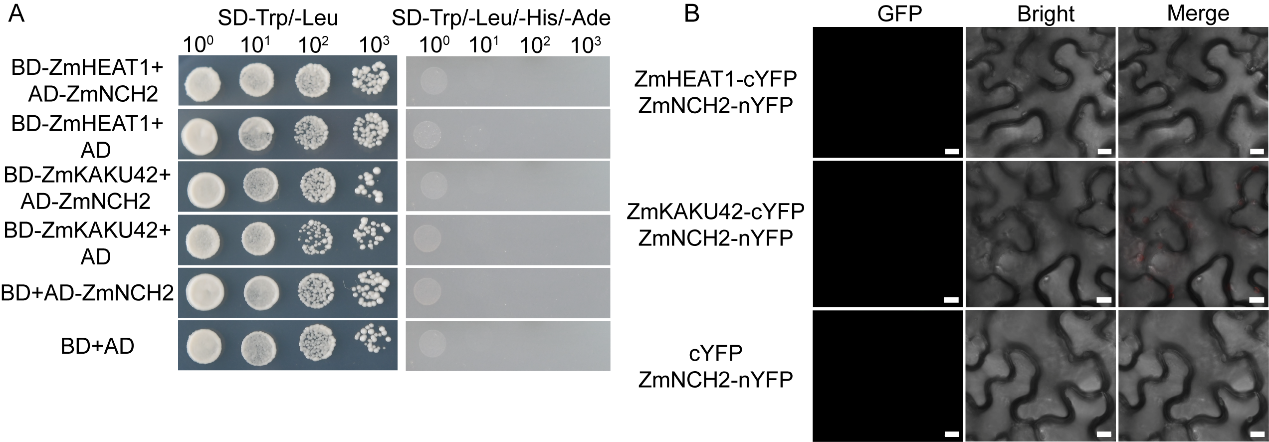

**Figure S19.** ZmHEAT1 and ZmKAKU42 do not interact with ZmNCH2.

A. Yeast two-hybrid assay showing that ZmHEAT1 and ZmKAKU42 do not interact with ZmNCH2. Yeast cells were grown on synthetic defined (SD) medium lacking Trp and Leu (−Trp −Leu) or Trp, Leu, His, and Ade (−Trp −Leu −His −Ade).
B. Bimolecular fluorescence complementation (BiFC) assay detecting no interaction between ZmHEAT1 or ZmKAKU42 and ZmNCH2. The YFP signal was visualized by confocal microscopy. Scale bars, 10 µm.

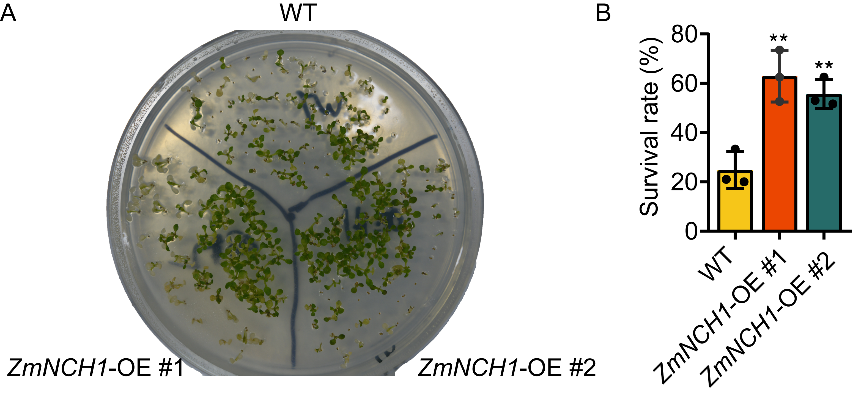

**Figure S20.** Effects of heat stress on seedling stage of the wild type and *ZmNCH1*-OE plants.

A. 7 days Arabidopsis seedlings were subjected to heat stress treatment in a 43°C water bath for 1.5 hour and recovery from HS for 7 days.
B. Survival rate of wild type and *ZmNCH1*-OE seedlings after recovery from HS for 7 days. Statistical significance was determined by one-way ANOVA; **, *p* < 0.01. Values are means ± SD with all individual data points shown as black dots.

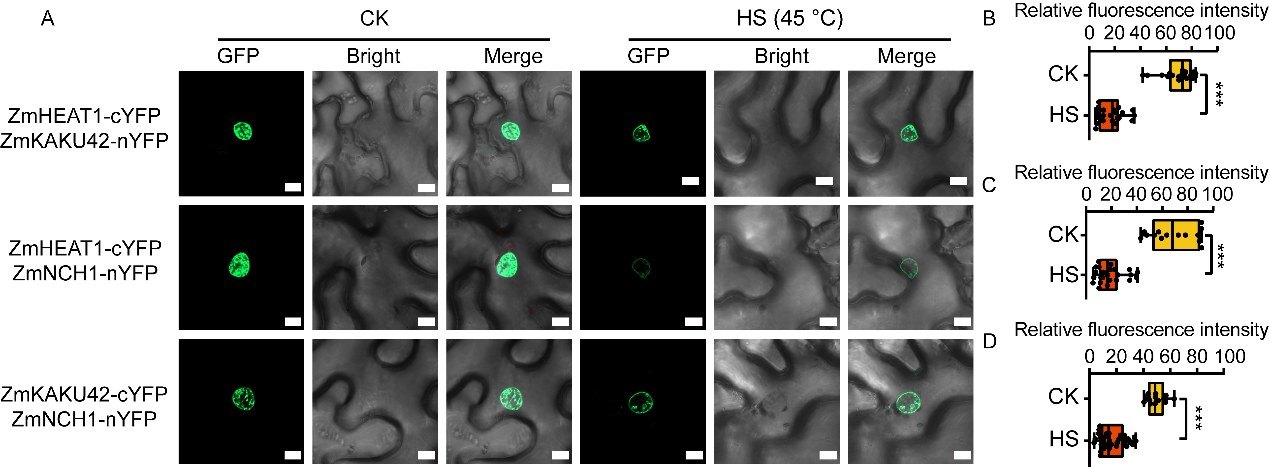

**Figure S21.** Changes in protein interaction intensity before and after heat stress treatment.

A. BiFC assay showing that ZmHEAT1 interact with ZmKAKU42 and ZmHEAT1, ZmKAKU42 interact with ZmNCH1 in *N. benthamiana* leaf cells. The YFP signal was visualized by confocal microscopy. Scale bars, 10 µm.

B-D. Interaction strength of proteins before and after heat treatment. Statistical significance was determined by one-way ANOVA; ***, *p* < 0.001. Values are means ± SD with all individual data points shown as black dots.

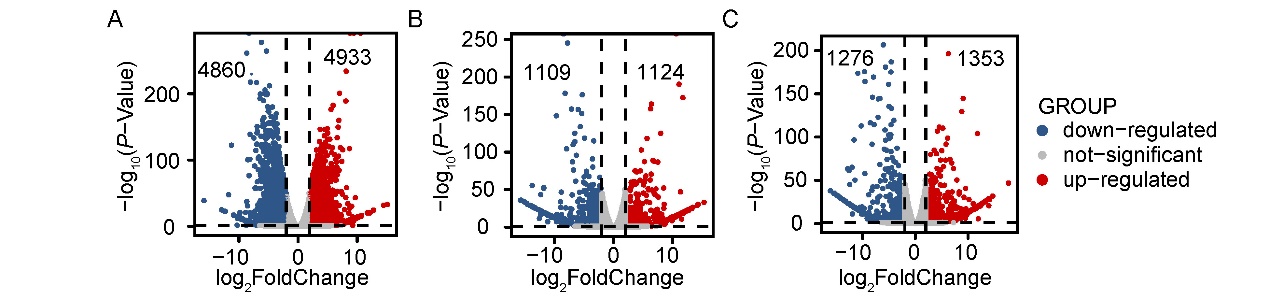

**Figure S22.** Transcriptome analysis of differentially expressed genes in the *Zmheat1* mutant relative to the wild type.

A–C. Volcano plots representing the fold-change in the expression levels of differentially expressed genes (DEGs) in the comparison groups of WT-0 h versus WT-45°C-24 h (A), *Zmheat1*-0 h versus WT-0 h (B), and *Zmheat1*-24 h versus WT-24 h (C) (*p* ＜0.05, absolute log_2_[fold-change]≥1.0). Gray dots represent genes with no significant change in expression. Three independent experiments were performed for each sample at each time point. The experiments were performed independently with three independent biological replicates (three biological replicates from different seedlings).

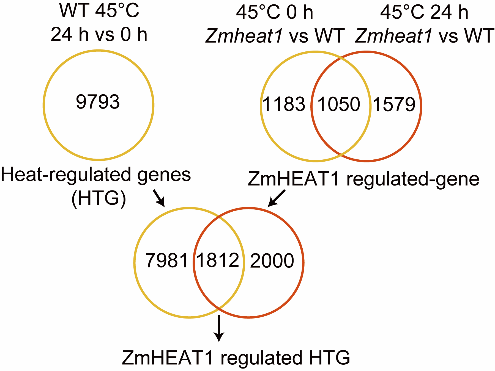

**Figure S23.** Venn diagrams showing the extent of overlap between differentially expressed genes (DEGs) regulated by heat (left), ZmHEAT1 (right), or both (middle).

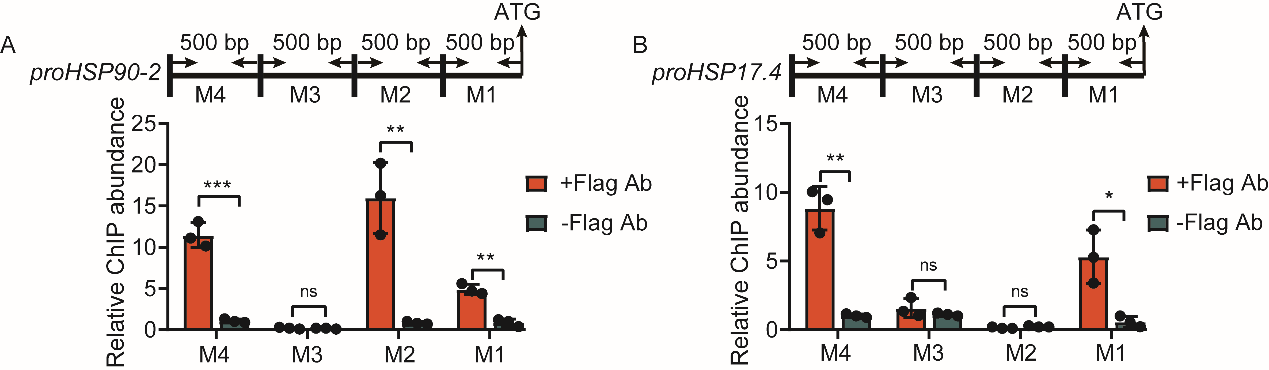

**Figure S24.** ChIP-qPCR analysis of ZmHEAT1 binding to four contiguous 500-bp fragments within the promoters of *HSP90-2* (*Zm00001d031332*) (A) and HSP17.4 (*Zm00001d039936*) (B). Chromatin was extracted from 14-d-old *ZmHEAT1*-OE transgenic seedlings and then precipitated with either an anti-flag antibody (+Ab) or only IgG (−Ab). Data are means ± SD (n = 3). Asterisks indicate significant differences determined by Student’s *t* test (ns, not significant, *, *p* < 0.05; **, *p* < 0.01, ***, *p* < 0.001).

Table S5. List of primers used in this study.

| Primer name | Primer sequences (5'-3') |
| --- | --- |
| ACTIN-F | CGATTGAGCATGGCATTGTCA |
| ACTIN-R | CCCACTAGCGTACAACGAA |
| qpcr-Zm00001d039936-F | AACGTGTTCGACCCCTTCTC |
| qpcr-Zm00001d039936-R | CAGTCTCGGAGTTGGTGGAG |
| qpcr-Zm00001d031332-F | CCAGCTGCTATCGCTCATCA |
| qpcr-Zm00001d031332-R | CCAAGGCATCGGAGGAGTTT |
| qocr-ZmHEAT1-F | GGACTGGGCGTAAAACCTGA |
| qocr-ZmHEAT1-R | AATGGGGATTGCCAAGGAGG |
| qPCR-NCH1-F | AGGTGGGTGCTACACGAAAG |
| qPCR-NCH1-R | TGAATGTGCTTCGCTCCCTT |
| At-actin-UBC2-F | TCAAATGGACCGCTCTTATC |
| At-actin-UBC2-R | CACAGACTGAAGCGTCCAAG |
| ZmHEAT1-GBKT7-F | ATGGAGGCCGAATTCATGGCGTCCCTCTTCGGGGC |
| ZmHEAT1-GBKT7-R | CAGGTCGACGGATCCCTACGTCGCTCTTCCTCTTG |
| ZmKAKU42-GADT7-F | GAGGCCAGTGAATTCATGGCGTCCCTCTTCGGGGC |
| ZmKAKU42-GADT7-R | GAGCTCGATGGATCCCTACGTCGCTCTTCCTTCTC |
| ZmKAKU42-GBKT7-F | ATGGAGGCCGAATTCATGGCGTCCCTCTTCGGGGC |
| ZmKAKU42-GBKT7-R | CAGGTCGACGGATCCCTACGTCGCTCTTCCTTCTC |
| ZmNCH1-GADT7-F | GAGGCCAGTGAATTCATGTTTACTCCGCAGGGCAA |
| ZmNCH1-GADT7-R | GAGCTCGATGGATCCTCATGTAGTGAAGAATGACC |
| ZmNCH2-GADT7-F | GAGGCCAGTGAATTCATGGCGAGCCCGCGCTCGGC |
| ZmNCH2-GADT7-R | GAGCTCGATGGATCCTCATGTGATGAGGAAACGCC |
| ZmHEAT1-flag-F | TCGACTCTAGAAAGCTTATGGCGTCCCTCTTCGGG |
| ZmHEAT1-flag-R | ATGGTACCGGATCCACTAGTCGTCGCTCTTCCTCTTGTTG |
| ZmHEAT1-GFP-F | CTAGAAAGCTTCTGCAGATGGCGTCCCTCTTCGGG |
| ZmHEAT1-GFP-R | CTTGCTCACCATGGTACCCGTCGCTCTTCCTCTTGTTG |
| ZmKAKU42-GFP-F | CTAGAAAGCTTCTGCAGATGGCGTCCCTCTTCGGGGC |
| ZmKAKU42-GFP-R | CTTGCTCACCATGGTACCCGTCGCTCTTCCTTCTCTCG |
| ZmNCH1-flag-F | TCGACTCTAGAAAGCTTATGTTTACTCCGCAGGGCA |
| ZmNCH1-flag-R | ATGGTACCGGATCCACTAGTTGTAGTGAAGAATGACCATA |
| ZmNCH1-GFP-F | CTAGAAAGCTTCTGCAGATGTTTACTCCGCAGGGCA |
| ZmNCH1-GFP-R | CTTGCTCACCATGGTACCTGTAGTGAAGAATGACCATA |
| ZmNCH2-GFP-F | CTAGAAAGCTTCTGCAGATGGCGAGCCCGCGCTCGGC |
| ZmNCH2-GFP-R | CTTGCTCACCATGGTACCTGTGATGAGGAAACGCCACA |
| ZmNCH1-MYC-F | TAAATACTAGTGGATCCGGTACATGTTTACTCCGCAGGGCAA |
| ZmNCH1-MYC-R | ATGAGCTTTTGCTCCATGGTACTCATGTAGTGAAGAATGACC |
| MBP-MYC-F | TAAATACTAGTGGATCCGGTACATGAATCACAAAGTGAGCTC |
| MBP-MYC-R | ATGAGCTTTTGCTCCATGGTACTCACGTTGGACCCGCGGATTGAG |
| MBP-flag-F | TCGACTCTAGAAAGCTTATGAATCACAAAGTGAGCTC |
| MBP-flag-R | ATGGTACCGGATCCACTAGTCGTTGGACCCGCGGATTGAG |
| ZmHEAT1-PGEX-4T-1-F | CGCGTGGATCCCCGGAATTCATGGCGTCCCTCTTCGGGGC |
| ZmHEAT1-PGEX-4T-1-R | TCACGATGCGGCCGCTCGAGCTACGTCGCTCTTCCTCTTG |
| ZmKAKU42-MBP-c5x-F | ATCGTCGACGGATCCATGGCCTCTTTGTTCGGTGC |
| ZmKAKU42-MBP-c5x-R | CGTTTTATTTGAAGCTTGGTGGCACGGCCTTCGC |
| ZmKAKU42-PGEX-4T-1-F | CGCGTGGATCCCCGGAATTCATGGCGTCCCTCTTCGGGGC |
| ZmKAKU42-PGEX-4T-1-R | TCACGATGCGGCCGCTCGAGCTACGTCGCTCTTCCTTCTC |
| ZmNCH1-PET32a-F | CCATGGCTGATATCGGATCCATGTTTACTCCGCAGGGCAA |
| ZmNCH1-PET32a-R | TGTCGACGGAGCTCGAATTCTGTAGTGAAGAATGACCATA |
| ZmHEAT1-SPYCE-F | GCCACTAGTGGATCCATGGCGTCCCTCTTCGGGGC |
| ZmHEAT1-SPYCE-R | AGCGGTACCCTCGAGCGTCGCTCTTCCTCTTGTTG |
| ZmKAKU42-SPYCE-F | GCCACTAGTGGATCCATGGCGTCCCTCTTCGGGGC |
| ZmKAKU42-SPYCE-R | AGCGGTACCCTCGAGCGTCGCTCTTCCTTCTCTCG |
| ZmKAKU42-SPYNE-F | GCCACTAGTGGATCCATGGCGTCCCTCTTCGGGGC |
| ZmKAKU42-SPYNE-R | AGCGGTACCCTCGAGCGTCGCTCTTCCTTCTCTCG |
| ZmNCH1-SPYNE-F | GCCACTAGTGGATCCATGTTTACTCCGCAGGGCAA |
| ZmNCH1-SPYNE-R | AGCGGTACCCTCGAGTGTAGTGAAGAATGACCATA |
| ZmNCH2-SPYNE-F | GCCACTAGTGGATCCATGGCGAGCCCGCGCTCGGC |
| ZmNCH2-SPYNE-R | AGCGGTACCCTCGAGTGTGATGAGGAAACGCCACA |
| ZmHEAT1-crispr-F | ATTGGTGGTTGAAAGAGC |
| ZmHEAT1-crispr-R | TGTCCTAATTTGAGCAGGT |
| ZmKAKU42-crispr-F | CACTGGTGCTCCTCCTCT |
| ZmKAKU42-crispr-R | CGAAACCTCCTGCCTAAT |
| ZmHEAT1-VM013-F | GACTCTAGAGGATCCATGGCGTCCCTCTTCGGGGC |
| ZmHEAT1-VM013-R | GTAGTCCATCCCGGGCGTCGCTCTTCCTCTTGTTG |
| ChIP-qPCR-Zm00001d031332-M1F | CGGCGTCGGATTTGGAAGGC |
| ChIP-qPCR-Zm00001d031332-M1R | GCCTTTGTGGCCTCCTCAAC |
| ChIP-qPCR-Zm00001d031332-M2F | TGGGTCTATTCTAACCCTTT |
| ChIP-qPCR-Zm00001d031332-M2R | GTCCAGTGCAGCTAGAAGAC |
| ChIP-qPCR-Zm00001d031332-M3F | AGTCCGGTGGCATACCGGAC |
| ChIP-qPCR-Zm00001d031332-M3R | ACTATATTAAAACATCAGTT |
| ChIP-qPCR-Zm00001d031332-M4F | CGAGACATGCTATAATGATG |
| ChIP-qPCR-Zm00001d031332-M4R | GTCGAGACGACCTTCCACGG |
| ChIP-qPCR-Zm00001d039936-M1F | AAATCCCTCAATTATAAGCG |
| ChIP-qPCR-Zm00001d039936-M1R | TCTAATGCGTTTAGTTGCAT |
| ChIP-qPCR-Zm00001d039936-M2F | AATAGTTTATGATGTATACT |
| ChIP-qPCR-Zm00001d039936-M2R | CACGGAAGGGGCGAGATGCG |
| ChIP-qPCR-Zm00001d039936-M3F | ACGTCGCTCGCTCGAACATC |
| ChIP-qPCR-Zm00001d039936-M3R | AAGCCCAGGCCAGACAAAGC |
| ChIP-qPCR-Zm00001d039936-M4F | CTAATAACTAAAGGAAGCCC |
| ChIP-qPCR-Zm00001d039936-M4R | TGTCGACTCTAGCTCTATGC |
